# Identifying parameters of host cell vulnerability during *Salmonella* infection by quantitative image analysis and modeling

**DOI:** 10.1101/139048

**Authors:** Jakub Voznica, Christophe Gardella, Ilia Belotserkovsky, Alexandre Dufour, Jost Enninga, Virginie Stévenin

## Abstract

*Salmonella* target and enter epithelial cells at permissive entry sites: some cells are more likely to be infected than others. However the parameters that lead to host cell heterogeneity are not known. Here, we quantitatively characterized host cell “vulnerability” towards *Salmonella* infection based on imaged parameters. We performed successive infections of the same host cell population followed by automated high-throuput microscopy and observed that infected cells have higher probability of being re-infected. Establishing a predictive model we identified two combined origins of host cell vulnerability: the pathogen-induced cellular vulnerability emerging from *Salmonella* uptake and persisting at later stage of the infection, and the host cell-inherent vulnerability. We linked the host cell inherent vulnerability with its morphological attributes such as the local cell crowding, and with host cell cholesterol content. This showed that the probability of *Salmonella* infection success can be forecast from morphological or molecular host cell parameters.

---

*Salmonella enterica* serovar Typhimurium (*S*. Typhimurium) is a Gram-negative bacterium that causes enteric diseases in many vertebrates after ingestion of contaminated food or water. Salmonellosis is one of the most common causes of food-borne diseases in humans and is considered to be major public health and global economic problem (26). After oral uptake, more than 99% of *S*. Typhimurium are killed in the stomach or in the gut (2). The surviving bacteria reach the distal ileum where they invade non-phagocytic intestinal epithelial cells (38). *In vitro* experiments have shown that *S*. Typhimurium invasion of host cells occurs after a phase of bacterial “Near Surface Swimming” (NSS) on the epithelial layer. The bacteria scan the surface and eventually stop and dock at a “selected” host cell (28, 37). Docking is irreversible (29) and followed by injection of *Salmonella* effectors into the host cell through a Type 3 Secretion System (T3SS), leading to the formation of ruffles that engulf the incoming bacterium (12, 22). Upon internalization *S*. Typhimurium either develops inside a *Salmonella*-Containing Vacuole (SCV) or it ruptures the SCV to escape into the cytoplasm where the pathogen replicates at a high rate, a phenomenon called hyperreplication (HR) (20, 33).

The mechanism by which *S*. Typhimurium targets specific host cellular sites for its entry remains debated. Santos and colleagues suggested that mitotic cells are selected due to increased cholesterol accumulation at the cell surface during metaphase (32). By contrast, Misselwitz and colleagues proposed that physical obstacles and forces that occur during the process of NSS lead to the targeting of topologically prominent sites, such as dividing cells or membrane ruffles (28). Finally, Lorkowski and colleagues have reported that the invasion of *S*. Typhimurium at the ruffle site is a highly cooperative effort (25, 29). Indeed, co-infection of WT and non-invasive *S*. Typhimurium mutants result in the entry of both strains inside the host cells: non-invasive *S*. Typhimurium mutants are trapped at ruffle sites and concomitantly internalized within the host cell, following active invasion by WT *S*. Typhimurium. However, the cooperative effect between intracellular and entering bacteria remains poorly understood at latter stage of the infection.

An increasing number of studies have highlighted the relevance of intrinsic cellular heterogeneity within eukaryotic monocultures. After seeding, cells display a dynamic range of variability in their morphology depending on their local microenvironment, including local density, and peripheral or central position within cellular islets (35). This heterogeneity results in differences of transcription (24, 9), lipid composition (9, 35) and sensitivity toward infections (35). Such cell-to-cell variations have been studied during viral infection revealing that simian virus 40 and mouse hepatitis virus present a population-determined pattern of infection associated with differential cell local crowding (35). In the context of bacterial infection, cell targeting has been related to bacterial cooperation at the entry site and evaluated at the whole population level using Colony Forming Unit (CFU) counting or flow cytometry analysis (25), but so far not *in situ* at the single cell level.

Here we investigated the susceptibility of epithelial host cells within the same cell population to become infected by *S*. Typhimurium. Our analysis revealed that some cells are more likely to be infected by *Salmonella* than others. We termed them “vulnerable cells”. The cell vulnerability was characterized in a quantitative manner by automated high-content imaging through double sequential infections with a delay of 1 to 3 h between the bacterial challenges. The number of intracellular bacteria per cell as well as the corresponding host cell parameters were assessed, such as cell perimeter, local density, and number of infected neighboring cells. Using a mathematical model, we showed that host cell vulnerability can be induced by a first bacterial uptake but also emerged from its intrinsic morphological and micro-environmental characteristics.

## RESULTS

### Sequential infections allow studies of *Salmonella* cooperation at the single cell level

We carried out a microscopy-based double infection assay to explore possible links between host cell vulnerability and successive bacterial infections of epithelial cells (Fig.1). HeLa cells grown in 96-well plates were subjected to a first infection with green *S*. Typhimurium expressing the fluorescent protein GFP (SL*_GFP_*) for 30 min followed by elimination of the extracellular bacteria via gentamicin treatment and washing. The cells were then incubated for 1, 2 or 3 h before being subjected to a second wave of infection with red *S*. Typhimurium expressing the fluorescent protein dsRed (SL*_dsRed_*). Extracellular bacteria were again eliminated in the same way, and the host cells were stained with CellMask and DAPI before automated image acquisition of entire culture wells (Fig.1A). The obtained images were analyzed with CellProfiler, a widely used image analysis software (1, 17) (Fig.1B). The differently labeled bacteria and the stained host cells enabled us to distinguish and quantify distinct cellular populations: those cells infected during the 1^st^ infection (I_1_) or not (noI_1_), those infected during the 2^nd^ infection (I_2_) or not (noI_2_), as well as the associated subpopulations (I_1_&I_2_, noI_1_&noI_2_, I_1_&noI_2_ and noI_1_&I_2_) (Fig.1C). We based our analysis on comparing probabilities of infection in these subpopulations.

**Fig. 1.**
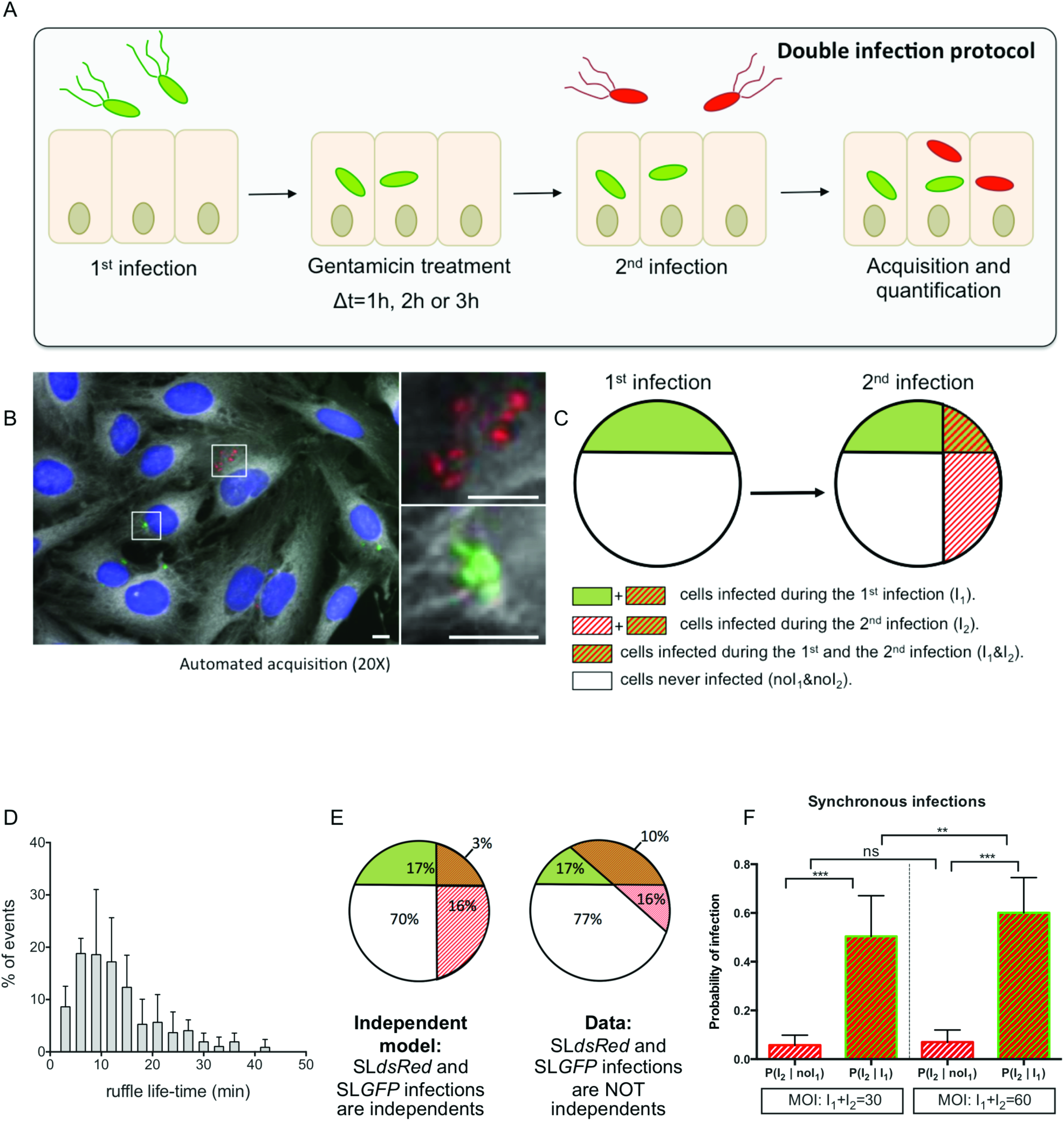
Double infections allow studies of *Salmonella* cooperation at the single cell level. **A. B. C.** Overview of the experimental workflow used in this study. **A.** Sequential infection protocol: HeLa cells grown in 96-wells plates since 24 h were subjected for 30 min to a first infection by SL*_GFP_*. This was followed by elimination of extracellular bacteria by gentamicin and incubation of the cells for 1, 2 or 3 h. The cells were subsequently challenged by a second infection with SL*_dsRed_* for 30 min. After removal of the extracellular bacteria, the samples were fixed. Nuclei were stained with DAPI and cell membranes were stained with CellMask before microscopic acquisition of the entire wells. **B.** Representative image of SL*_GFP_* and SL*_dsRed_* internalized in HeLa cells. Host cell nuclei are visible through DAPI (in blue), and cell membranes through CellMask (in grey). Scale bar correspond to 5μm. **C.** Scheme of our statistical analysis of different subpopulations. The following cellular populations can be distinguished: those cells infected during the 1^st^ infection (I_1_) or not (noI_1_), those infected during the 2^nd^ infection (I_2_) or not (noI_2_), along with the related subpopulations (I_1_&I_2_, noI_1_&noI_2_). This scheme maps the case of two independent infections. **D.** Time distribution of the ruffle disappearance during *Salmonella* infection followed in actin-GFP transfected cells by time-lapse microscopy. **E.** Comparison of an independent model (left) with the obtained data (right). The percentages are averaged from 6 independent experiments, represented in **C** with an MOI of 30. **F.** Comparison of the conditional probability of infection for two different populations during synchronous infection of SL*_GFP_* and SL*_dsRed_*. The MOIs were calculated after averaged CFU counting from 6 different experiments. P-values were obtained after paired t-test.

### Cooperation at the entry site during the presence of ruffles

In order to test the reliability of our method, we analyzed first if we could reproduce previously published results on the ruffle-dependent cooperation between individual salmonellae during host cell entry (28, 25). To do this we determined first the time window during which ruffle-associated cooperation could potentially occur performing time-lapse microscopy of *Salmonella* infection of HeLa cells (Fig. 1D) and Caco-2 cells (data not shown) transiently expressing GFP-tagged actin and labeled with the membrane dye FM 4-64 respectively. Time series of 90 min at 3 min intervals provided image sequences with forming and disappearing ruffles. In most of the cases, we observed for both cell lines the uptake of one to two bacteria per ruffle, and we saw ruffle disappearance in less than 15 min (Video.S1). We noticed that the more bacteria were engulfed by the ruffles, the longer we could detect the presence of these ruffles. Therefore, newly arriving bacteria prompted additional growth of the ruffles (Video.S2). We quantified the ruffle lifetime by measuring the delay of their disappearance after the entry of the last bacterium. The few cases of very high infection (>5 bacteria/ruffle) that could not be properly analyzed were excluded. Quantification revealed an average ruffle lifetime of 13 min and that 90% of the ruffles completely disappeared after 24 min (Fig.1D). The results for Caco-2 cell infection were similar to those of HeLa cells.

We then challenged HeLa cells with SL*_GFP_* and SL*_dsRed_* at the same time and compared the probability for SL*_dsRed_* to infect the same cell containing simultaneously SL*_GFP_* with those that did not contain SL*_GFP_* (Fig.1E and Fig.1F); *see Materials and Methods for details*. The repartition of the different populations of infected cells (Fig.1E) shows a much larger overlap between the cells co-infected with SL*_GFP_* and SL*_dsRed_* than one would anticipate theoretically for two independent infections. Thus, the efficiency of *Salmonella* invasion in an individual epithelial cell depends on the concomitant invasion of the same cell by other salmonellae. The difference of the concomitant infection probability (Fig.1F) was striking as it was 8 times more likely for SL*_dsRed_* to infect a cell also infected by SL*_GFP_* than a cell not infected by SL*_GFP_*. Interestingly, increasing the multiplicity of infection (MOI) resulted in a significant increase of the SL*_dsRed_* infection in cells infected by SL*_GFP_*, but not in cells not infected by SL*_GFP_*. This result confirmed that the direct effect of an MOI increase is a higher number of bacteria that infect certain cells rather than an increase of the overall number of cells that become infected. It underlines the relevance of ruffle-associated cooperation between salmonellae at the entry site. Taken together, these results validated that our system was operational.

### The probability of being re-infected by *Salmonella* is higher for already-infected cells, even after the disappearance of the entry ruffles

To study long-term and ruffle-unrelated cooperative events of *Salmonella* co-infections, we set the sequential infections with a delay of 1 h between the two infection waves, killing extracellular bacteria in between through gentamicin treatment. Scanning our time-lapse movies, we were ensured that this time lag led to the complete absence of any remaining entry ruffles from the first infection. In addition, we extended the delay between the two sequential infections to 2 h and 3 h (*see* Fig.1A). We compared the different populations of cells infected during the 2^nd^ infection (population I2), depending on whether they were already infected during the first wave of infection (population I_2_ | I_1_) or not (population I_2_ | noI_1_) for HeLa (Fig.2A) and Caco-2 (Fig.2B) cells. For both tested cell types, it was significantly more probable for a cell infected the 1^st^ time to be re-infected the 2^nd^ time compared to a cell not previously infected. We propose that such cells are somehow more vulnerable for future infection.

**Fig. 2.**
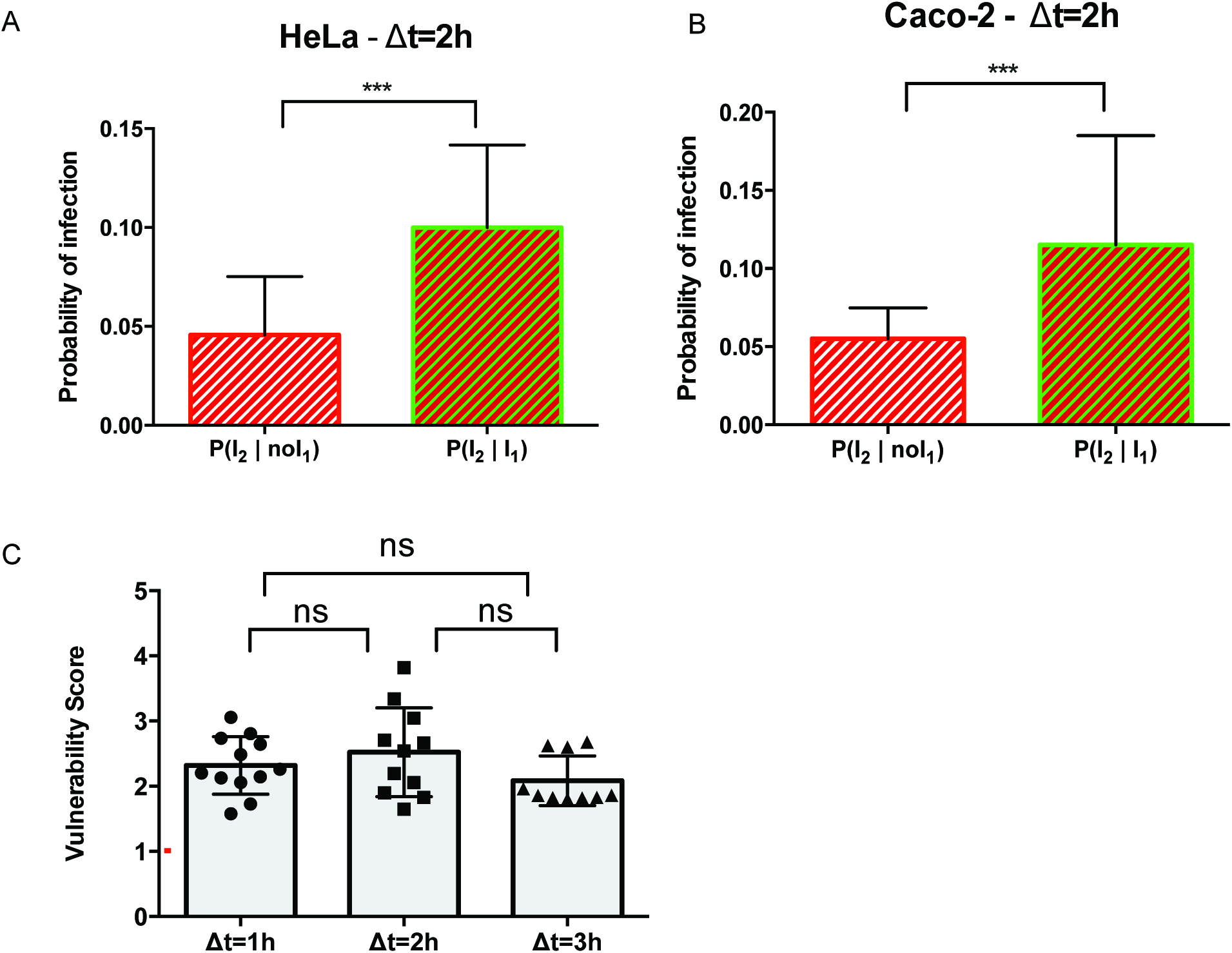
The probability of being re-infected by *Salmonella* remains higher for already infected cells aCer entry ruffle disappearance. **A-B.** Conditional probability of infection for two different populations during sequential infection with a delay of 2 h for HeLa cells (**A**) and Caco-2 cells (**B**). Results were obtained from 3 independent experiments and P-values were obtained after paired t-test. **C.** The vulnerability score was plobed for infection with a 1, 2 or 3 h delay before the second infection. The red line corresponds to P(I_2_| I_1_)=P(I_2_ | noI_1_) = 1 indicating the independence of the infections I_2_ and I_1_. Values above the red line correspond to P(I_2_ | I_1_) >P(I_2_ | noI_1_) indicating a cooperation between infections. Values below the red line correspond to P(I_2_ | I_1_) <P(I_2_ | noI_1_) indicating a competition between infections. Results were obtained from 3 independent experiments per time-point, and P-values were obtained after unpaired t-test.

During all sequential infection experiments we also controlled the overall infection efficiencies of SL*_GFP_* and SL*_dsRed_* at all measured time points (1^st^: SL*_GFP_* - 2^nd^: SL*_dsRed_* or in the reverse order) (**Fig.S1**). In all cases, the percentage of cells infected by each fluorescent *Salmonella* was similar for cells subjected to single (control) or sequential infections, underlining that sequential infections did not change the overall infection efficiency for the differently colored salmonellae. Nevertheless, we noticed a decrease of the amount of infected cells between the early infection and later time points. This effect is most likely due to the technically unavoidable gentamicin treatment between infections. Besides, SL*_GFP_* showed a higher infectivity than SL*_dsRed_* for each condition explained by general deleterious effects of the heterologously overexpressed fluorescent proteins on *Salmonella* infectivity, and by the partial loss of dsRed expression observed by us and others. Taking into account these issues, we took advantage of the observed consistency of the differences of infection efficiency between the initial and the successive infections, and between SL*_GFP_* and SL*_dsRed_*. This consistency allows comparative analyses of the ratio of the different infection probabilities, and it provided us with an analytical tool for precise quantification independently of the variances of the differently colored bacteria and technical hurdles of sequential infection.

We defined a “vulnerability score” as the conditional probability for a cell to be infected during the 2^nd^ infection after it had been already infected during the 1^st^ one (I_2_ | I_1_), divided by the conditional probability for a cell to be infected during the 2^nd^ infection when it had not been previously infected (I_2_ | noI_1_) (described in more detail in *Materials and Methods*). We also analyzed changes of the vulnerability score in time comparing cells subjected to sequential infection with 1, 2 and 3 h delays (Fig.2B and **Fig.S2** for detailed representation of the conditional probability for each replicate). Surprisingly, the vulnerability score appeared un-altered. We obtained similar results inverting the order of the used pathogen, infecting first with SL*_dsRed_* and then with SL*_GFP_* (**Fig.S3**). It was not possible to shorten the delay between infections to less than 1 h due to the ruffle influence, and we could not extend it beyond 3 h due to potential release of hyper-replicative (HR) bacteria from the first infection into the extracellular medium that could then re-infect new cells during the 2^nd^ wave of infection. Altogether, these results showed that, after ruffle disappearance, the infected cells remain more vulnerable to a new infection than the non-infected ones, and this vulnerability is stable in time.

### Cell vulnerability to secondary infection can be predicted from the number of intracellular bacteria

So far, we only considered the character “infected” or “non-infected” for each cell after SL*_GFP_* and SL*_dsRed_* infections that provides global trends on their interaction. To further exploit our data we quantified the number of bacteria per host cell and related the obtained numbers with the previously extracted vulnerability scores. The distribution of intracellular bacteria inside infected cells at 2.5 h post-infection (pi) showed that most of the cells contained few bacteria, and the proportion of cells with higher number of intracellular bacteria decreases drastically. Overall, we were able to distinguish three groups of infected cells: the ones containing one to two intracellular bacteria (35% of the global population), the ones containing three to eight intracellular bacteria (39% of the global population) and the ones containing more than nine intracellular bacteria (26% of the global population), corresponding respectively to low, medium and high infections (Fig.3A).

**Fig. 3.**
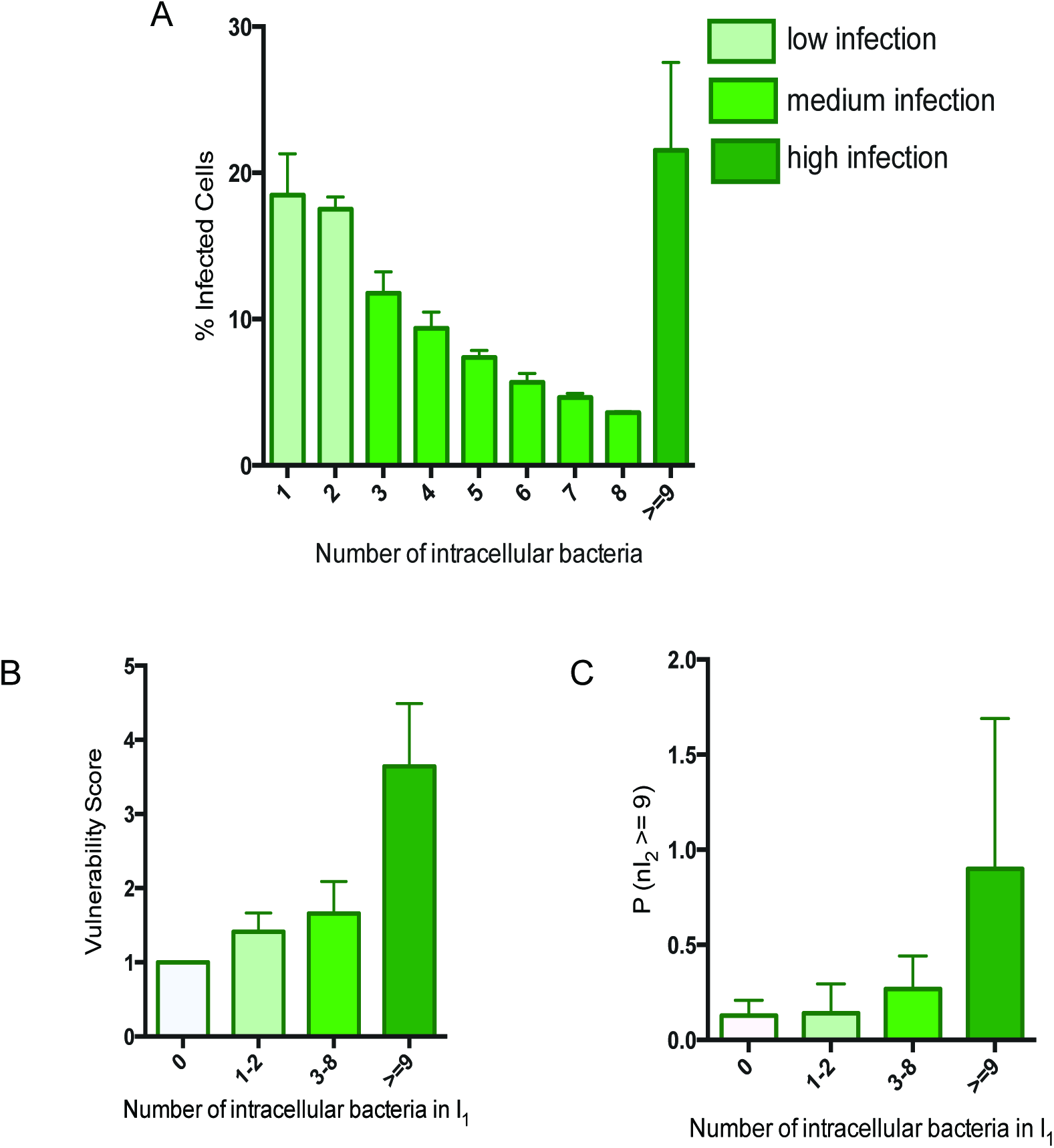
Cell vulnerability can be predicted from the number of bacteria previously internalized. **A.** Distribution of the number of intracellular bacteria detected at 1.5 h pi (average from 3 replicates). The infection efficiencies are clustered in 3 groups: low, medium and high infection, corresponding respectively to 1 to 2; 3 to 8 or more than 9 bacteria per cell. **B.** The vulnerability score is represented as a function of the number of intracellular bacteria resulting from the 1^st^ infection. **C.** Probability of a cell to be highly infected during the 2^nd^ infection (nI_2_ ≥9) as a function of the number of intracellular bacteria being internalized during the 1^st^ infection. **B** and **C** represent the data merged from all the experiments (delay of 1, 2 and 3 h before the second infection). Groups of infection efficiency are identical in **A**, **B** and **C**.

We compared the vulnerability score of these three infection groups during sequential double infections (Fig.3B). This analysis revealed that the more bacteria had entered in a given host cell during the first infection, the more it was likely that this cell became re-infected. Such tendencies still emerged when the bacteria were not grouped, but analyzed individually, underlining the robustness of this result (**Fig.S4**).

Then, we investigated how the level of bacterial uptake during the second infection depends on the number of intracellular bacteria of the first infection. For this we quantified the probability for a cell to be highly infected during the second infection as a function of the efficiency of the first uptake (Fig.3C). We found that the more intracellular bacteria had been internalized during the first infection, the more likely they were to ingest a high amount of new bacteria during the second infection. Therefore, we propose that cell vulnerability is maintained from the first to the second infection.

### Cell vulnerability as intrinsic or induced property

The results from the sequential infections (Fig.3 and Fig.4) provided quantitative scores of cell vulnerability towards *Salmonella* infection. We secondly investigated the origin of the observed cell vulnerability. Two possibilities can be anticipated: (i) the cellular vulnerability would be an intrinsic host cell attribute (hypothesis 1: “intrinsic vulnerability”) or (ii) it would be induced by bacterial uptake (hypothesis 2: “induced vulnerability”) (Fig.4A). In theory, these hypotheses can be distinguish by the observable different probability of the 2^nd^ wave of infection occurring in previously non-infected cells P(I_2_ | noI_1_) as depicted in the two schemes of Fig.4B and described as follows: In the case of vulnerability as intrinsic attribute, the probability of infection P(I_2_ | noI_1_) would be lower than P(I_2Ctr_) as the pool of vulnerable cells would be already partially consumed during the 1^st^ sequential infection, whereas it would remain conserved in the control (Fig.4B**-left**). In the case of induced vulnerability, the probability of infection P(I_2_ | noI_1_) would be similar to P(I_2Ctr_), as the cells would be considered with equivalent vulnerabilities before their first infection (Fig.4B**-right**). The experimental data obtained did not show a significant difference between P(I_2_ | noI_1_) and P(I_2Ctr_) (t-test p-value >0,05) (Fig.4C), suggesting that vulnerability may be induced by bacterial uptake (Fig.4B, *hypothesis 2*). Taking into account the small percentage of cells belonging to the studied subpopulations we caution that the absence of a statistically significant difference between these populations did not allow to exclude the first hypothesis of host cell inherent vulnerability.

**Fig. 4.**
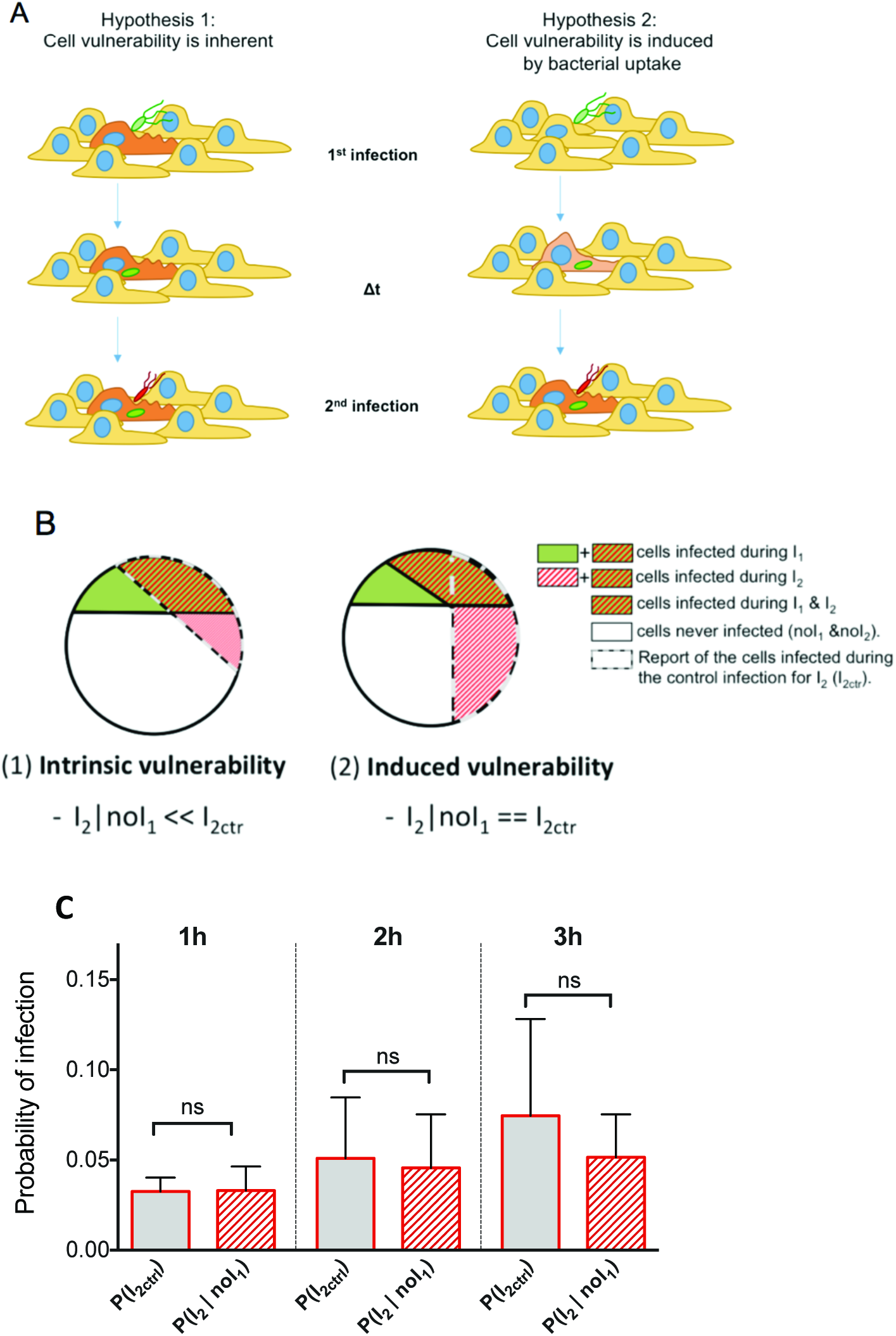
Cell vulnerability examined as an intrinsic or an induced property. **A.** Schemes of the two hypotheses for the origin of cell vulnerability. In the hypothesis 1, cell vulnerability is inherent: some cells (in orange) are more vulnerable towards infection than other cells (in yellow). In the hypothesis 2, cell vulnerability is induced by bacterial uptake: before infection cells are equal regarding their vulnerability (in yellow), but after infection the infected cells turn progressively more vulnerable (in orange). **B.** Graphic representation of the theoretical distribution of the different populations in the case of hypothesis 1 (left) or hypothesis 2 (right). **C.** Probability of infection during sequential infection with 1, 2 and 3 h delays for control cells (I_2Ctr_) and cells non infected during the 1^st^ infection (noI_1_). P-values were obtained after unpaired t-test (P(I_2ctr_) vs P(I_2_ | noI_1_)).

### Single cell vulnerability to *Salmonella* infection is a combination of intrinsic and induced vulnerability

Considering that the subpopulation comparison could not exclude an involvement of inherent vulnerability, we developed a mathematical model to evaluate the relative contribution of induced and inherent vulnerability to the overall cell vulnerability towards *Salmonella* infection. To investigate the contribution of cell parameters at the single-cell level, we measured different intrinsic variables that could influence cellular vulnerability, namely the cell morphology (cell perimeters, cell circularity), the local environment (local cell density, number of infected and non-infected neighboring cells), and the above-analyzed features of the *Salmonella* infection (delay between infections, load of intracellular bacteria per cell from I_1_) (Fig.5A). We extracted all these elements using Icy, an image analysis software (6) recently used for *Salmonella* infection studies *in situ* (27) (*see* **Fig.S5A** for illustration of Icy cell segmentation).

**Fig. 5.**
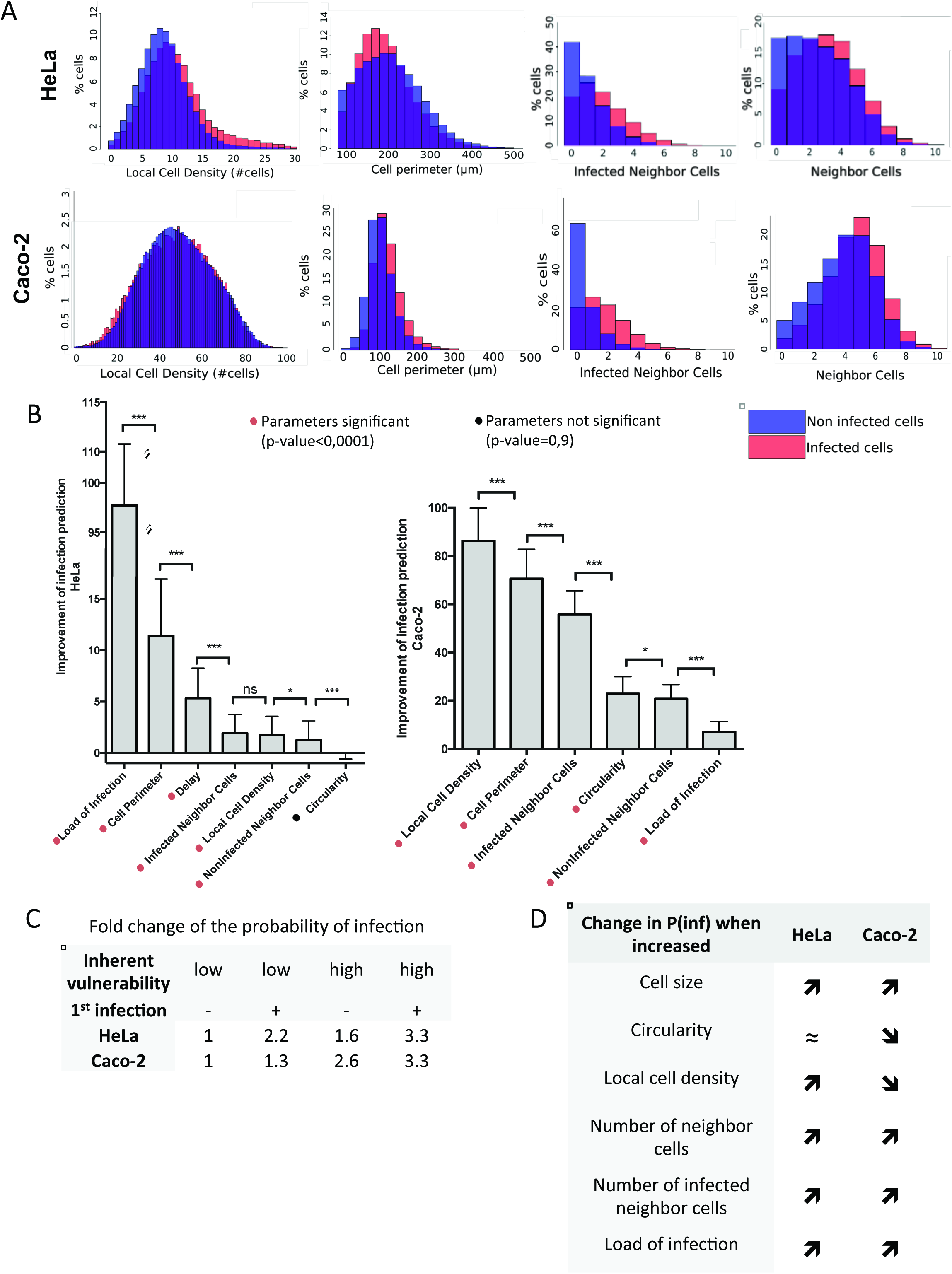
Single cell vulnerability to *Salmonella* infec0on is a combina0on of intrinsic and induced vulnerability. **A.** The depicted cellular parameters were determined for HeLa cells (upper panel) and Caco-2 cells (lower panel) as described in detail in *Materials and Methods*. An overlay of the distribution of some of these parameters in infected (red) or non-infected (blue) cells is shown. **B.** Quantification of the improvement of infection prediction by each cell parameters by subtracting the likelihood (in log) of the model including all parameters from a model ignoring one parameter. Results are averaged over 100 training/testing circles for each model. P-values were obtained after paired t-test. **C.** Fold change of the probability of infection as a function of the intrinsic vulnerability and of a previous infection. **D.** Increasing or decreasing of the probability of infection when the listed cell parameters increase their values.

First, we analyzed the distribution of distinct cellular parameters in either infected or in non-infected HeLa (upper panels) and Caco-2 (lower panels) cell populations (Fig.5A). For both cell types, the infected cells displayed distinct cellular features in comparison to the non-infected cells, such as a higher local crowding reflected by a higher number of neighboring cells in direct contact. Comparing the relative correlations of the cellular parameters, we highlight the presence of strong links between a number of them (**Fig.S5B-C, S6** and **S7**). In particular cell morphology is highly dependent on the local micro-environment, such as the local cell density that negatively correlates with the cell perimeter in HeLa and Caco-2 cells. Interestingly, cells infected during the second bacterial challenge are more likely to be nearby cells infected during the first bacterial challenge (“infected neighbor cells”) than by non-infected neighbor cells. Thus *Salmonella* infection of one cell increases the probability of its neighboring cells to be subsequently infected.

To quantify the direct involvement of each studied parameter on the overall cell vulnerability we developed a statistic modeling approach adapted to our high-throughput microscopy dataset on sequential *Salmonella* infection. This model is based on a logistic regression able to predict the infection efficiency at the single cell level from cellular parameters. We measured the contribution of each parameter for the prediction by estimating how well the model predicts compared to a model that would ignore one parameter; as described in *Materials and Methods* (Fig.5B). Taken separately, the load of intracellular bacteria resulting from I_1_ directly improved the prediction of cell vulnerability towards subsequent infection (Fig.5B). Thus, host cell vulnerability is induced by bacterial uptake, which is in line with our experimental data. In addition, the host cell parameters linked with cell morphology and local environment also significantly improved the model prediction of infection for HeLa and for Caco-2 cells (*see* **Table1** and **Table2** *for model details and the value of the coefficients*). Together, our modeling approach revealed that single host cell vulnerability to *Salmonella* infection is a combination of intrinsic and bacterial-induced vulnerability.

We quantified their relative involvement by calculating the model-based fold change of the probability of infection for a cell not infected and having a low score of inherent vulnerability with a cell infected and/or having a high score of inherent vulnerability (Fig.5C). This showed that induced and intrinsic vulnerability have both a strong impact on the overall cell vulnerability. Interestingly, the induced vulnerability is more prevalent for *Salmonella* infection of HeLa cells (2,2 fold-increase) than infection of Caco-2 cells (1,3 fold-increase), whereas the inherent vulnerability plays a more prominent role for Caco-2 cell infections (2,6 fold-increase) than for HeLa cells (1,6 fold-increase). From these findings we conclude that the analyzed host cell parameters are differentially involved in relation to cell vulnerability towards *Salmonella* infection depending of the cell type. In particular, the local cell density increases the cell vulnerability for HeLa cells but decreases it for Caco-2 cells (Fig.5D). This could be explained by the polarization of the Caco-2 at high confluency and highlights the specificity of each predicted model for a given cell-type.

We also investigated whether the first infection affects the inherent host cell parameters, we compared the correlation between parameters that were identified as being either involved or not involved in the inherent vulnerability of the cell. As their correlations were similar in infected and non-infected cells (data not shown) we concluded that *Salmonella* infection did not impact the implication of the studied inherent cell parameters.

### Reliability of the model-based prediction of infection

To investigate the spatial distribution of the cell vulnerability among the cell population, we created “vulnerability maps” from original images of the cell population after labeling each cell nucleus with a color corresponding to its probability of infection (Fig.6A). Notably, we could confirm that on average the infected cells were properly assigned with a higher prediction score to be infected than the non-infected ones (*see **Fig.S8** for quantification*). Based on our vulnerability maps, the predicted infected cells showed a very good overlap or were in close vicinity with the experimentally infected cells (Fig.6A). This illustrates the reliability of our approach in a qualitative way, and it also underlines the impact of local micro-environment on cell vulnerability. We went on and quantified the veracity of the HeLa and Caco-2 adapted models when confronted with 100 experimentally measured infected and 100 experimentally measured non-infected cells. For both cell-types, models allowed a good prediction in the majority of the cases, 62% for HeLa and 66% for Caco-2 respectively (Fig.6B). Taken together, these results attest that the probability of *Salmonella* infection success can be forecast at the near single-cell level based on host cell parameters.

**Fig. 6.**
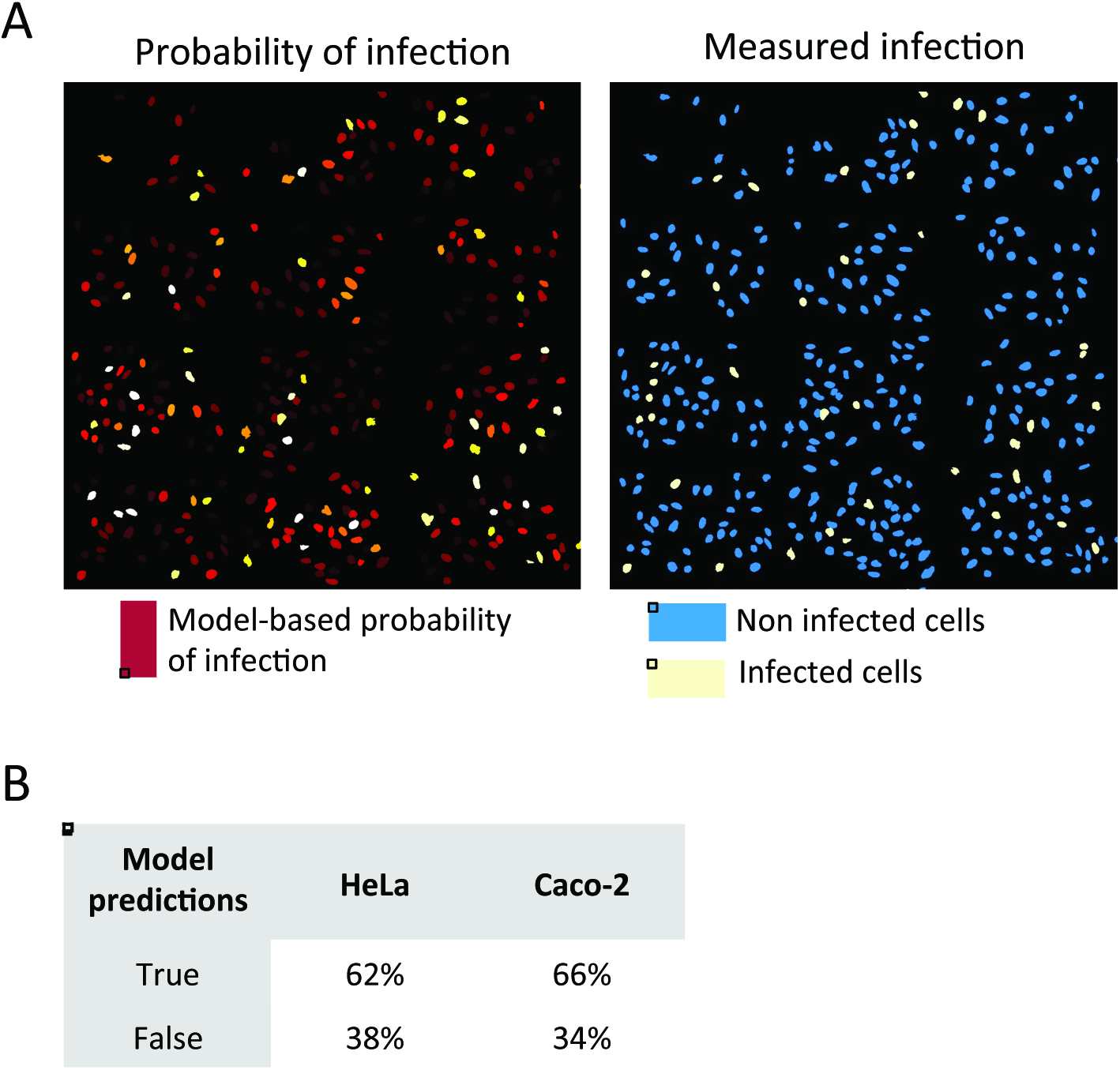
Comparison of model-predicted vulnerability of single-cell with measured-infection. **A.** Model-predicted probability of infection displayed on reproduced original images of HeLa cells (left panel). Colors are adapted for maximum contrast between lowest (deep red) and highest (white) probability of infection. Measured infections from experiments are shown (top-right panel). **B.** Estimation of the reliability of the two (HeLa and Caco-2) models developed when tested on a total of 100 infected cells and 100 non-infected cells.

### Involvement of cellular cholesterol rate as an inherent vulnerability factor

To investigate molecular players that are linked with the inherent cell vulnerability to *Salmonella* infection, we analyzed the plasma membrane composition as main feature known to be relevant for *Salmonella* infection. We focused on cholesterol as cells at low crowding present a higher amount of free cholesterol than cells at high crowding (9). We monitored the relation between global cellular cholesterol levels and host cell targeting performing *Salmonella* infection of HeLa cells for 30 min followed by cholesterol labeling via filipin staining and flow cytometry analysis (Fig.7). For each experiment, we binned the total cell population into five subpopulations corresponding to increasing cellular levels of cholesterol that we classified as 1 to 5, with each subpopulation containing 20% of the total cells (*see **Fig.S9** for FACS gating details*). Comparing the number of infected cells in these different subpopulations with different amounts of cholesterol, we revealed that the probability of infection decreased with increasing cholesterol levels. From this we conclude that cells with a lower amount of cholesterol are preferentially targeted by *Salmonella* compared to those with higher cholesterol levels.

**Fig. 7.**
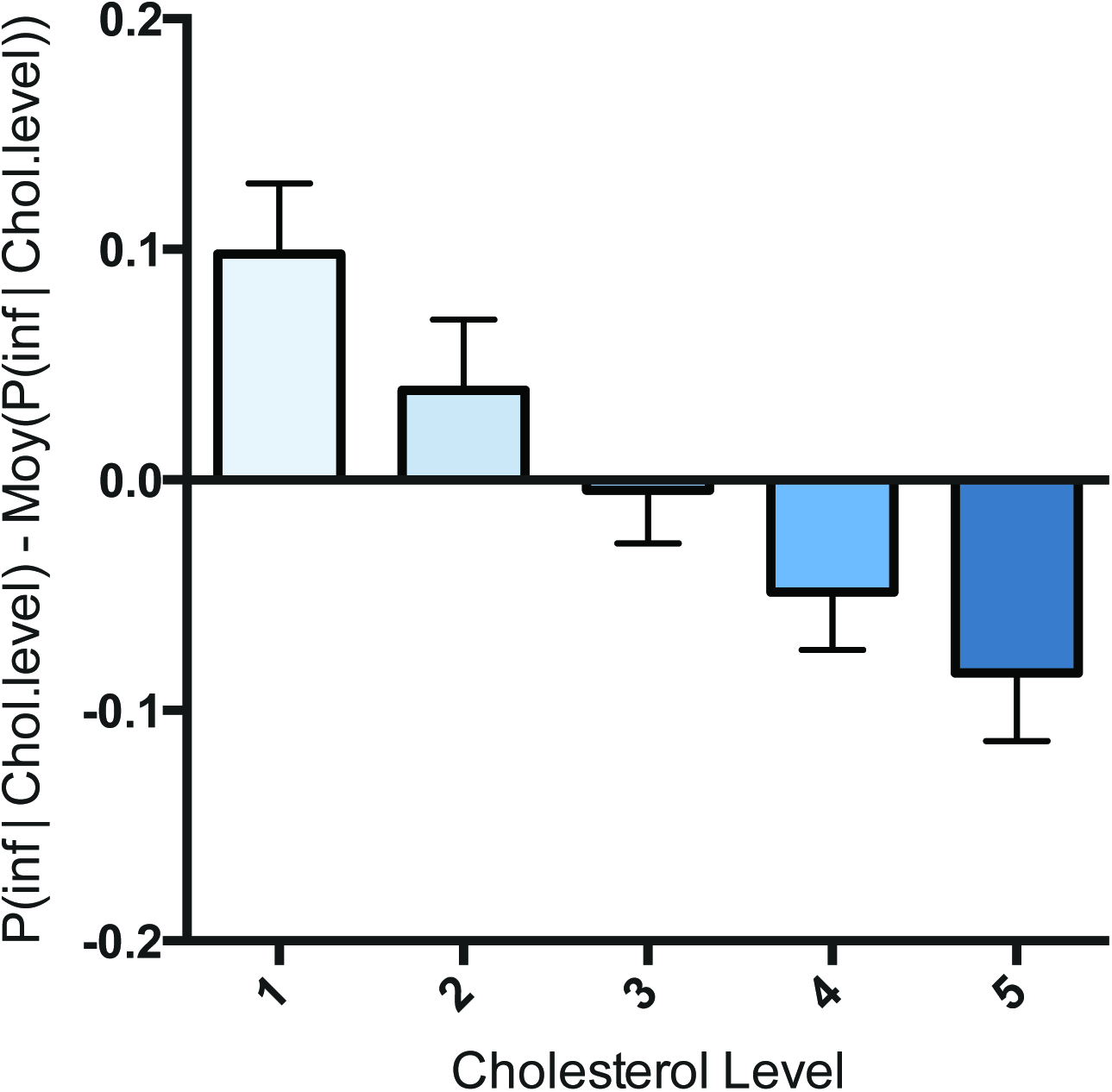
Probability of infection as a function of single cell cholesterol level. Variation of the probability of *Salmonella* infection at different levels of host cholesterol measured by FACS as described in detail in the text. Cholesterol levels were binned in five categories at 20% steps from lowest to highest levels over the total cell population, each category contains 20% of the whole cells. Results are averaged over 3 independent experiments.

## DISCUSSION

Cellular heterogeneity describes cases in which genetically identical cells present different behaviors and morphologies. This biological phenomenon is commonly present in an epithelial layer of an individual as well as within a monolayer of cultured cells. Despite the realization of the importance of cellular heterogeneity, its study has only become feasible during recent years, mainly thanks to the implementation of novel technologies such as imaging and computer-assisted analysis. In the context of pathogen infection, this heterogeneity produces cells unequally vulnerable or resistant which impacts on the overall infection.

We investigated the cell vulnerability of epithelial cells for *S*. Typhimurium infection. According to our results, infected cells display a strikingly higher probability of being re-infected with *Salmonella*, even after disappearance of membrane ruffles. We obtained similar results in two relevant epithelial cell lines, HeLa and Caco-2, suggesting that this represents a conserved propensity towards *Salmonella* infection. The measured cellular vulnerability remained unaltered for all measured time-points ranging from a delay of 1 h to 3 h between the infections. Attributing a “vulnerability score” to the challenged cells, we showed a higher vulnerability score in cells previously infected, and we found that this score increases with the amount of intracellular bacteria contained by a given cell. This result raises the issue of the bacterial impact on the cell vulnerability. Therefore, we aimed at distinguishing inherent cell vulnerability from the one induced by bacterial uptake (Fig.4A, *hypothesis 1 and 2 respectively*) exploiting the imaging data obtained via a high-content analytical pipeline. This allowed visualization of the infection *in situ* and provided a large number of associated cellular parameters. We quantified the implication of specific parameters associated with individual cells on the cell vulnerability towards *Salmonella* infection. It appeared clearly that the efficiency of early bacterial uptake during the first infection directly determines cell vulnerability. Thus *Salmonella* induce an increase of the cell vulnerability toward subsequent infections.

While long-term cooperation between bacteria has been intensively studied for communities of bacteria living in a common extracellular environment (5), little is known about the cooperation between intracellular and extracellular bacteria leading to increased bacterial uptake. Nevertheless, this phenomenon has been investigated more extensively for many viruses, including bacteriophages (4), influenza virus (14), poxviruses (7, 21), flaviviruses (39, 34), alphaviruses (18), and alphaherpesviruses (3). Generally, those works have demonstrated that the first virus to infect a cell has the capacity to prevent co-infection of other viruses belonging either to the same strain, or to more distantly related or unrelated strains. It is termed “superinfection exclusion” and may protect limited cellular resources and promote the replication and dissemination of the originally infecting virus. By analogy, the increased probability of cellular re-infection by *Salmonella* can be phrased as a “superinfection promotion”. It remains to be clarified if such process is relevant for all intracellular bacteria. For instance and in contrast to *Salmonella* infection, Jorgensen *et al* reported that the *Chlamydia* effector protein CPAF secreted from bacteria within mature inclusions prevents those that are still extracellular to invade (16). Thus, CPAF could be a factor mediating *Chlamydia* resistance towards superinfection.

Our approach also allowed the relative quantification of the impact of different host cell parameters on the inherent vulnerability of host cell to *Salmonella* infection. In particular, morphological attributes and local cell crowding are highly linked with this vulnerability. Cell crowding as major determinant for the probability to become infected has been proposed by Snijder and colleagues in the context of viral infection. They showed that during infections by the simian virus SV40 or the mouse hepatitis virus (MHV), the targeted cells have different localization within cell islets (35). SV40 and MHV infect preferentially either peripheral or central cells, a phenomenon that is linked with the differential expression levels of focal adhesion kinase and the presence of sphingolipid GM1 at the plasma membrane of challenged host cells. Thus, similarly to several viral infections, the probability of infection of a single cell by *Salmonella* is influenced by its local environment.

Our analytical tools will be useful for further studies on *Salmonella*, and for other researchers working on different intracellular bacteria pathogens, such as *Chlamydia*, *Listeria* or *Shigella* (*see Materials and Methods*). We revealed that some cells are indeed intrinsically more vulnerable to *Salmonella* and will be targeted by the bacteria first. Most of the tested parameters appeared relevant for model-based infection prediction but are differentially involved in the cell vulnerability depending on the cell-type studied. Developing an adapted model based on host cell parameters we could forecast the probability of *Salmonella* infection success at the near single-cell level. Interestingly, the number of infected neighboring cells is highly increased in the population of infected cells. Cases of bacterial uptake impacting on the cells neighboring the infection (called bystander cells) have been previously reported for *Shigella* that induces an IL-8 immune response after NFκB activation detectable from 2 h pi in 70% of the bystander cells (19). However, it is not known whether the neighboring cells are also more sensitive towards *Shigella* entry.

Because of our incomplete knowledge of host factors that are involved in the early attachment, such as potential entry receptors, it remains difficult to identify the molecular mechanisms that establish the differential vulnerability during *Salmonella* infection. Although receptors for direct recognition of *Salmonella* have been proposed, such as the cystic fibrosis transmembrane conductance regulator (CFTR) (31) and the epithelium growth factor receptor (EGFR) (30), many cell types infected by *Salmonella* do not express them (15). Therefore, it has been proposed that recognition mechanisms likely involve more ubiquitous factors (10). To explore molecular cues involved in the inherent heterogeneity of host cell vulnerability, we decided to investigate the membrane lipid composition, in particular cellular cholesterol. This was based on previous studies showing that the amount of free cholesterol per cell negatively correlates with the local cell crowding (9). We found that an increase of cholesterol amount at single cell level correlates with a lower vulnerability of the cell, so that *Salmonella* preferential target host cells with low amounts of cholesterol. Intriguingly, these results on an implicated host molecule are in agreement with the morphological feature of local density; HeLa cells at high density contain lower amounts of cholesterol and display an increased inherent cell vulnerability. The role of cholesterol during *Salmonella* infection is still debated. Several studies have demonstrated that the *Salmonella* SipB effector and translocon component requires cholesterol for proper functioning (13, 10). In this context, it should be noted that the translocons operate in small cholesterol-rich microdomains at the plasma membrane and cannot be linked readily with the overall cholesterol levels. Furthermore, those studies were based on sterol sequestering agents and biosynthesis inhibitors. Contrastingly, Gilk and colleagues showed that cholesterol is not essential for *Salmonella* invasion and intracellular replication inside host cells using an original mouse model, (11). In our study we highlighted that non-treated cells with a low amount of global cellular cholesterol are preferentially targeted by *Salmonella*, which does not exclude a potential involvement of cholesterol at the subcellular level.

In conclusion, our study represents a first step in understanding *Salmonella* cell targeting and provides a path for the identification of cellular and bacterial factors involved in host cell vulnerability. Such factors could be targeted to render a cell more resistant to pathogen infections, allowing potential new therapeutic strategies. Together, our study delineates in a quantitative manner the importance of vulnerable cell recognition and bacterial cooperation for cell targeting by *S*. Typhimurium.

## MATERIALS AND METHODS

### Bacterial Strains

The following *S.* Typhimurium were used: SL1344 (wild type), SL1344 pM965 (*Salmonella*-GFP) described by Stecher *et al* (36), and SL1344 pGG2 (*Salmonella-*dsRed) obtained after transformation of SL1344 with the pGG2 plasmid described by Lelouard *et al* (23). Bacteria were grown in Lysogeny Broth (LB) medium supplemented with 0.3 M NaCl and ampicillin at 50 µg/ml at 37°C in an orbital shaker.

### Cell Culture

All cell culture reagents were purchased from Invitrogen unless otherwise stated. Human epithelial HeLa cells (clone CCL-2 from ATCC) were cultured in Dulbecco's Modified Eagle's Medium (DMEM) supplemented with 10% (v/v) fetal bovine serum (FBS), at 37 °C, 5% CO_2_. Intestinal epithelial Caco-2 TC7 cells (kindly provided by P. Sansonsetti) were grown in DMEM supplemented with 10% FBS at 37°C, 10% CO_2_. All infection assays were performed in EM buffer (120 mM NaCl, 7 mM KCl, 1.8 mM CaCl_2_, 0.8 mM MgCl_2_, 5 mM glucose, 25 mM HEPES, pH 7.4). HeLa cells were transfected with pEGFP-actin plasmid DNA (8) from a maxiprep, using the X-tremeGENE 9 DNA transfection reagent (Roche) for 48 h.

### Double Infection Assays

For invasion experiments, overnight bacterial cultures were sub-cultured 1/20 and grown until late exponential/early stationary phase. Before infection, bacteria were gently washed and resuspended in EM buffer. Bacteria were added to the cells at an MOI of 30 corresponding to CFU, and incubated for 30 min at 37 °C, 5% CO_2_. Non-internalized bacteria were eliminated by washing 3 times with warm EM buffer and incubated for 1, 2 or 3 h at 37 °C, 5% CO_2_. Adding EM buffer containing 100 µg/ml gentamicin for 1 h killed extracellular bacteria. The concentration of gentamicin was then decreased to 10 µg/ml and 10% FBS was added to the medium. At the desired time points, the cells were washed again in EM buffer to eliminate the remaining gentamicin and re-infected with a fresh batch of sub-cultured bacteria following the same protocol. After killing the extracellular bacteria again by a 1 h of incubation with EM buffer containing 100 µg/ml gentamicin, the cells were fixed with 4% paraformaldehyde at room temperature for immunofluorescence analysis.

### Microscopy

All image acquisitions were performed on a Nikon inverted widefield microscope using a 20x/0.5NA air objective, an automatic programmable XY-stage and the Nikon perfect focus system. For sequential infections, 161 fields were imaged per well and four channels per field were captured using a CoolSnap2 camera (Roeper Scientific). Nuclei and cells were stained using DAPI (excitation/emission wavelengths: 350/470 nm) and the cell bodies with CellMask DeepRed Plasma Membrane Stain (ThermoFisherScientific, excitation/emission wavelengths: 640/670 nm) respectively. Caco-2 cells were stained with the FM® 4-64 membrane dye (Invitrogen) before time lapse imaging (excitation/emission wavelengths: 558/734 nm). Quantification of the ruffle timing was performed on the same microscope, using a 20×/0.5NA air objective and time intervals of 3 min for 90 min. Time lapse imaging of ruffles was performed on a DeltaVision widefield microscope using a 60×/1.42 NA oil objective and z-stacks with a spacing of 500 nm. The images were subsequently deconvolved using DeltaVision Elite integrated software.

### Cholesterol measurements

HeLa cells were challenged with SL*_GFP_* for 30 min before trypsinization, fixation with 4% paraformaldehyde and incubation with 16ug/mL filipin complex from *Streptomyces filipinensis* (Sigma-Aldrich). This treatment was directly followed by FACS measurement on *BD FACS CANTO* cytometer using the excitation/emission wavelengths of 405/450 nm and 488/530 nm for filipin and GFP fluorescence detection respectively. Infected and non-infected cells were distinguished using the green fluorescence emitted by SL*_GFP_* (*see **Fig.S9** for gating details*). Data were processed using FlowJo software.

### Image Analysis

All images were analyzed with two open source software: CellProfiler (http://cellprofiler.org/) and Icy (http://icy.bioimageanalysis.org/). CellProfiler was used to detect each single cell and the number of its intracellular salmonellae expressing either GFP or dsRed. The following modules were used during the analysis: *IdentifyPrimaryObjects* recognized nuclei and bacteria; *IdentifySecondaryObjects* identified cells (here the secondary objects) by extending the nuclear area previously recognized; *RelateObjects* assigned bacteria within individual cells. Icy was used for accurate detection of cell borders and the cellular microenvironment analysis. We used a graphical environment called *Protocols* for the development of an analytical pipeline including the following plugins: *HK*-*Means* that identify nuclei by pre-filtering the signal to identify objects within a size range; *Spot Detector* that identify bacteria; *Active Contours* that identify the edges of the plasma membrane by propagating the Region of Interest (ROI) detected for the nuclei; and *Javascript* that parent the ROI of cells with bacteria, to measure local cell density and to distinguish which neighboring cells are infected by which bacteria.

### Probability

P(I_2_|I_1_) means “Probability of the 2^nd^ sequential infection, knowing that the cell has been infected by the 1^st^ one” and is calculated as follows:

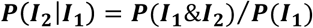

Where *P(I_1_)* = [*Number of cells in I_1_ / Total number of cells*], and *P(I_1_&I_2_)* = [*Number of cells in I_1_&I_2_ / Total number of cells*].

P(I_2_|noI_1_) means “Probability of the 2^nd^ sequential infection, knowing that the cell has not been infected by the 1^st^ one” and is calculated as follows:

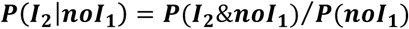

Where *P(noI_1_)* = [*Number of cells in noI_1_ / Total number of cells*], and *P(I_2_&noI_1_)* = [*Number of cells in noI_1_&I_2_ / Total number of cells*].

### Model

We modeled the influence of multiple parameters on the probability of a second infection. A Boolean variable Y represents the second infection: It is equal to 1 for infected cells and 0 otherwise. Its probability is predicted by the following seven parameters: *Load of infection* (LOI) represents the number of infecting bacterium during the first infection, separated in 4 groups corresponding to no (0 bacteria), low (1 or 2), medium (3 to 8) or high (9+) infection. *Delay* is a categorical variable corresponding to the delay between the 1^st^ and the 2^nd^ infections (1, 2 or 3 h). *Infected neighbor cells* (X1) refers to the number of cells in contact that had been infected during the first infection. *Non-Infected neighbor cells* (X2) refers to the number of cells in contact which had not been infected during the first infection. *Local Cell Density* (X3) is the number of cells present in a vicinity of 100 µm. The distance is calculated between the center of the nuclei. *Cell perimeter* (X4) is the length of the perimeter of the cell (in µm) obtained after segmentation. *Circularity* (X5) refers to the cell circularity defined as: “4π*area/perimeter^2^”. This parameter is higher for circular cells, and lower for cells that are elongated or have complex shape, but does not depend *a priori* on the cell size. In practice we used to its square root. The probability of Y during the second infection is modeled as:

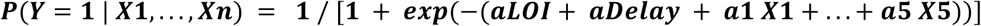

where aLOI (resp. aDelay) has a different value for each of the LOI categories (resp. Delay categories), and a_1_,…,a_5_ are constants. All parameters were learned by maximizing the likelihood of the model, e.g. the probability of the observed data as measured by the model. We used 115 000 and 327 000 cells to train and test the model for HeLa and Caco-2 cells respectively. We divided the cell population into two random sets; the training set (9/10^th^ of the cells per replicate) and testing set (1/10^th^ of the cells) and computed the likelihood of infection observed in the testing set. The higher the likelihood, the better the parameters of the model predicted infection. We repeated this procedure 100 times. To measure the improvement of infection prediction by taking into account each parameter, the likelihood of the complete model was compared (on a log scale) with the likelihood of seven models ignoring each time one parameter. This difference of log-likelihood is reported in Fig.5B.

Quantification of the impact of a parameter towards cell vulnerability was obtained by applying our statistical model to the 1^st^ and the 3^rd^ quantile values of a given parameter, while other parameters were kept equal at their median values. We obtained the probabilities of the second infection for these two sets and reported their ratio. In Fig.5D, the arrows “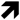” and “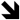” correspond to a ratio above and under 1 respectively. The parameters-values corresponding to a low inherent vulnerability of HeLa and Caco-2 cells were the following: local cell density (1^st^ quantile and 3^rd^ quantile respectively), cell perimeter (1^st^ quantile), infected neighboring cells (median), non-infected neighboring cells (median), circularity (median and 3^rd^ quantile respectively). The parameters-values corresponding to a high inherent vulnerability of HeLa and Caco-2 cells were the following: local cell density (3^rd^ quantile and 1^st^ quantile respectively), cell perimeter (3^rd^ quantile), infected neighboring cells (median), non-infected neighboring cells (median), circularity (median and 1^st^ quantile respectively).

Models reliability was evaluated using 100 infected and 100 non-infected cells and quantifying the amount of “good predictions” among those cells. We repeated this procedure 100 times and showed the average. As a comparison, a random model would provide approximately 50% of “good predictions”.

### Statistical analysis

The statistical analysis was performed using R and GraphPad Prism. T-tests were used to evaluate the significance of the results, referred like *, **, *** for p*-*values <0.05, <0.01, and <0.001, respectively.

## Data availability

The pipeline used on CellProfiler and on ICY, as well as the R code used to generate the model can be provided by the authors.

## Acknowledgments

We thank Jennifer Fredlund and Andrew Rutenberg for their help during the initial phase of the project, Adrien Sauvaget, Claude Loverdo, Kristine Schauer and Uriel Hazan for productive discussions, and all the members of the DIHP unit and BioImage Analysis Group for helpful interactions. VS was supported by a Ph.D. fellowship from the University Paris Diderot attributed by the ENS Cachan, Université Paris-Saclay. JE is member of the LabEx consortia IBEID and MilieuInterieur. JE also acknowledges support of from the ANR (grant StopBugEntry and AutoHostPath) and the ERC (CoG EndoSubvert).

## Author contributions

VS and JE designed the study and initiated the project. VS, JE and AD supervised the project. *Salmonella* infection were conducted by JV and VS. The automatic image analysis method was designed by VS, JV and AD. Flow cytometry measurements were achieved by IB and VS. The statistical analysis method was conceived by CG, JV and VS and performed by JV. VS and JE wrote the manuscript. JE provided the funding.

---

**Vldeo.S1.**
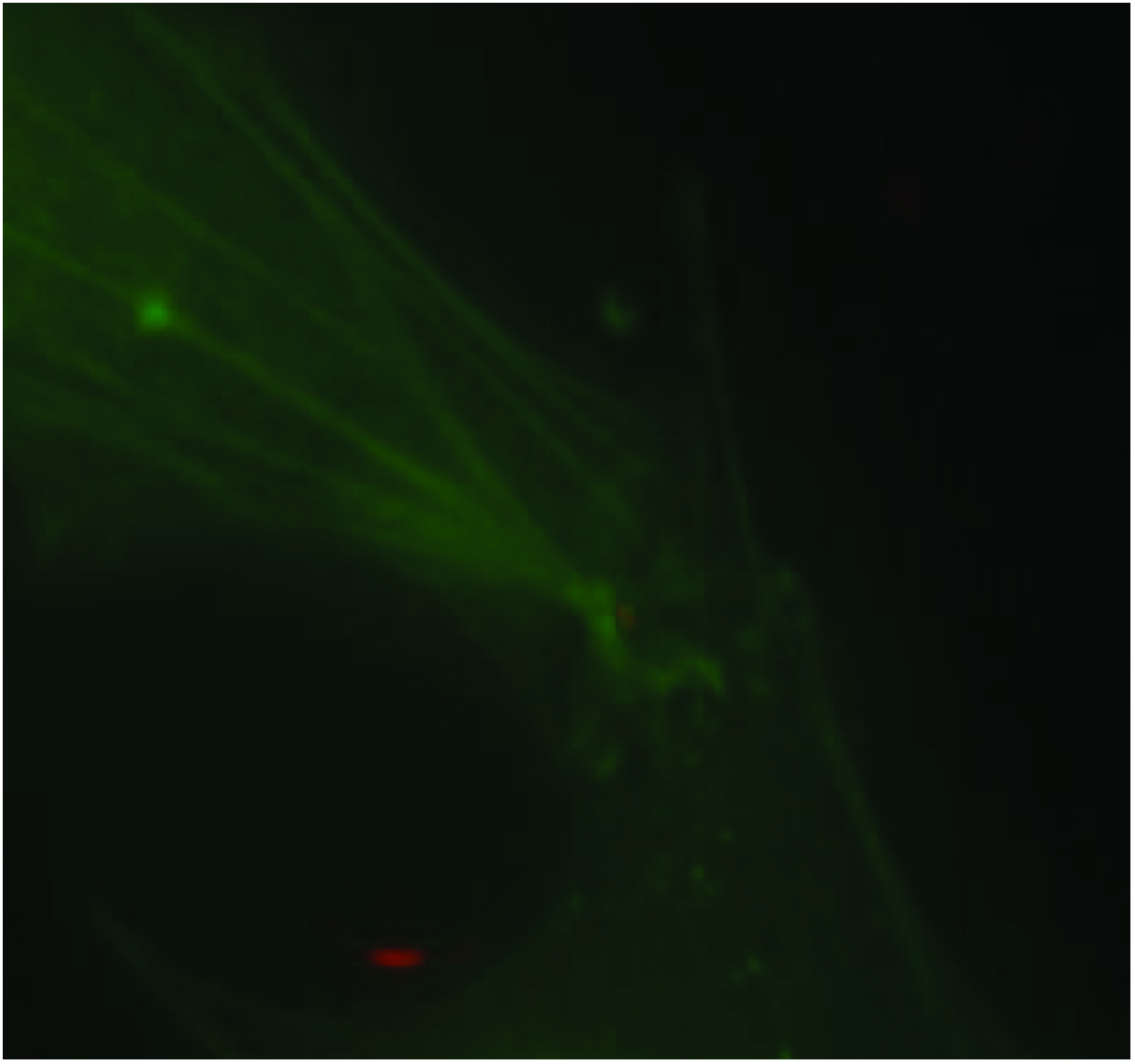
Ruffle appearance and disappearance after entry of single salmonellae in a host cell. Time intervals between the frames are 3 min. The green channel corresponds to actin-GFI > transfected cells and shows the membrane ruffles. The red channel shows salmonellae SL*_dsRed_*.

**Video.S2.**
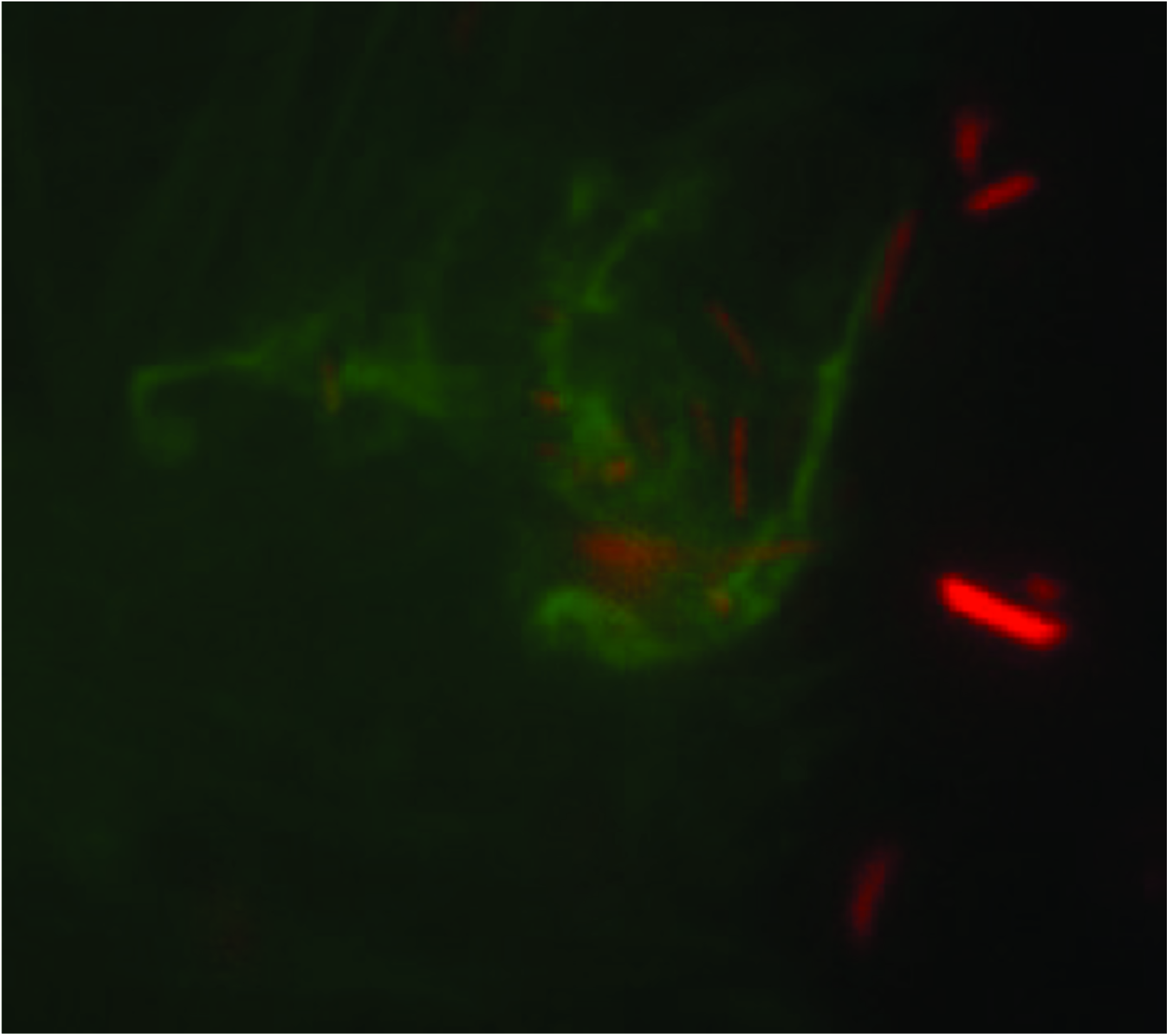
Ruffle appearance and disappearance after entry of multiple salmonellae in a host cell. Time intervals between the frames are 3 min. The green channel corresponds to actin-GFP transfected cells and shows the membrane ruffles. The red channel shows salmonellae SL*_dsRed_*.

**Table S1.**
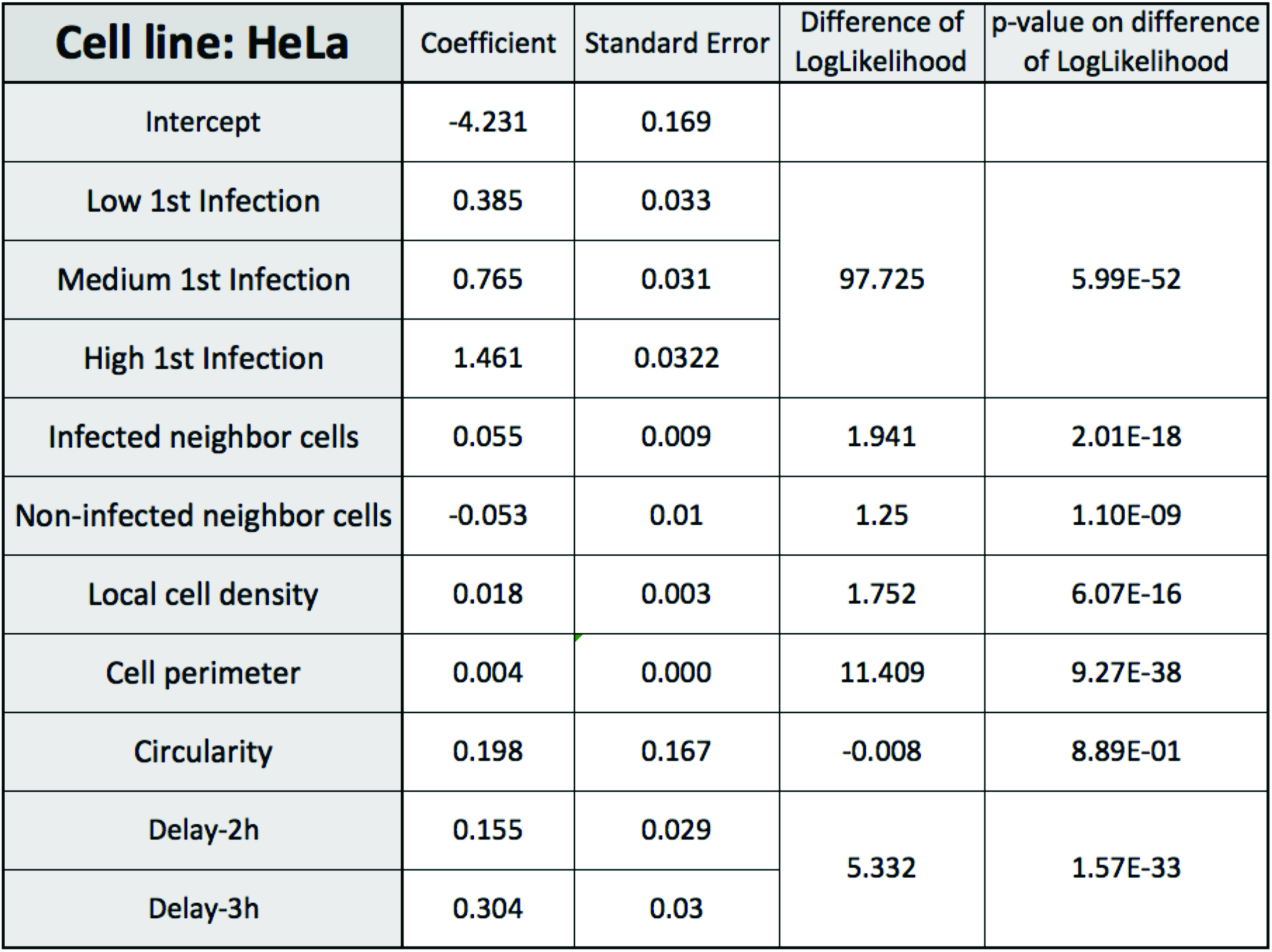
Model coefficient values for Hela cells with the corresponding standard error for each cell parameter. Difference of log-likelihood obtained aOer subtrac;on of the loglikelihood of the model including all parameters from the log-likelihood of a model ignoring one parameter (see graphic representation in Fig.5B). The presented values were averaged with the values obtained over 100 training/testing circles for each model. P-values were obtained after paired t-test.

| **Cell line: HeLa** | Coefficient | Standard Error | Difference of LogLikelihood | p-value on difference of LogLikelihood |
| --- | --- | --- | --- | --- |
| Intercept | -4.231 | 0.169 |  |  |
| Low 1st Infection | 0.385 | 0.033 | 97.725 | 5.99E-52 |
| Medium 1st Infection | 0.765 | 0.031 |  |  |
| High 1st Infection | 1.461 | 0.0322 |  |  |
| Infected neighbor cells | 0.055 | 0.009 | 1.941 | 2.01E-18 |
| Non-infected neighbor cells | -0.053 | 0.01 | 1.25 | 1.10E-09 |
| Local cell density | 0.018 | 0.003 | 1.752 | 6.07E-16 |
| Cell perimeter | 0.004 | 0.000 | 11.409 | 9.27E-38 |
| Circularity | 0.198 | 0.167 | -0.008 | 8.89E-01 |
| Delay-2h | 0.155 | 0.029 | 5.332 | 1.57E-33 |
| Delay-3h | 0.304 | 0.03 |  |  |

**Table S2.** Model coefficient values for Caco-2 cells with the corresponding standard error for each cell parameter. Difference of log-likelihood obtained after subtrac;on of the loglikelihood of the model including all parameters from the log-likelihood of a model ignoring one parameter (see graphic representation in Fig.5B). The presented values were averaged with the values obtained over 100 training/testing circles for each model. P-values were obtained after paired t-test.

| **Cell line: Caco-2** | Coefficient | Standard Error | Difference of LogLikelihood | p-value on difference of LogLikelihood |
| --- | --- | --- | --- | --- |
| Intercept | -0.48 | 0.14 |  |  |
| Low 1st Infection | 0.25 | 0.02 | 7.07 | 2.29E-30 |
| Medium 1st Infection | 0.23 | 0.03 |  |  |
| High 1st Infection | 0.20 | 0.06 |  |  |
| Infected neighbor cells | 0.23 | 0.01 | 55.69 | 3.28E-77 |
| Non-infected neighbor cells | 0.09 | 0.00 | 20.71 | 7.75E-58 |
| Local cell density | -0.02 | 0.00 | 86.25 | 6.70E-82 |
| Cell perimeter | 0.01 | 0.00 | 70.56 | 2.52E-78 |
| Circularity | -2.96 | 0.14 | 22.83 | 1.26E-53 |

**Fig. S1.**
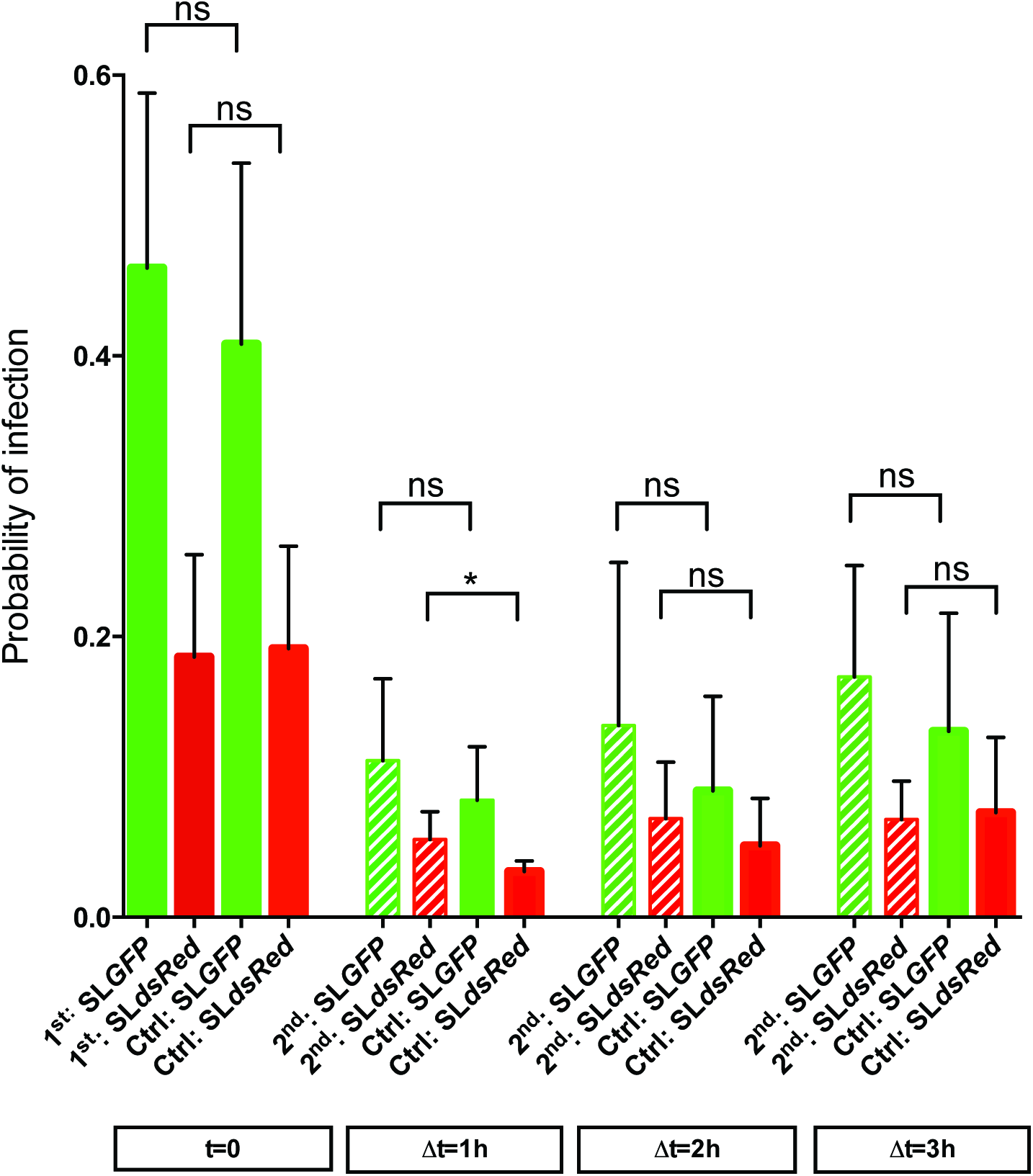
Probability of SL*_GFP_* and SL*_dsRed_* infections at different time-points after the beginning of cell challenge (t=0) between single (control) or sequential infections.

**Fig. S2.**
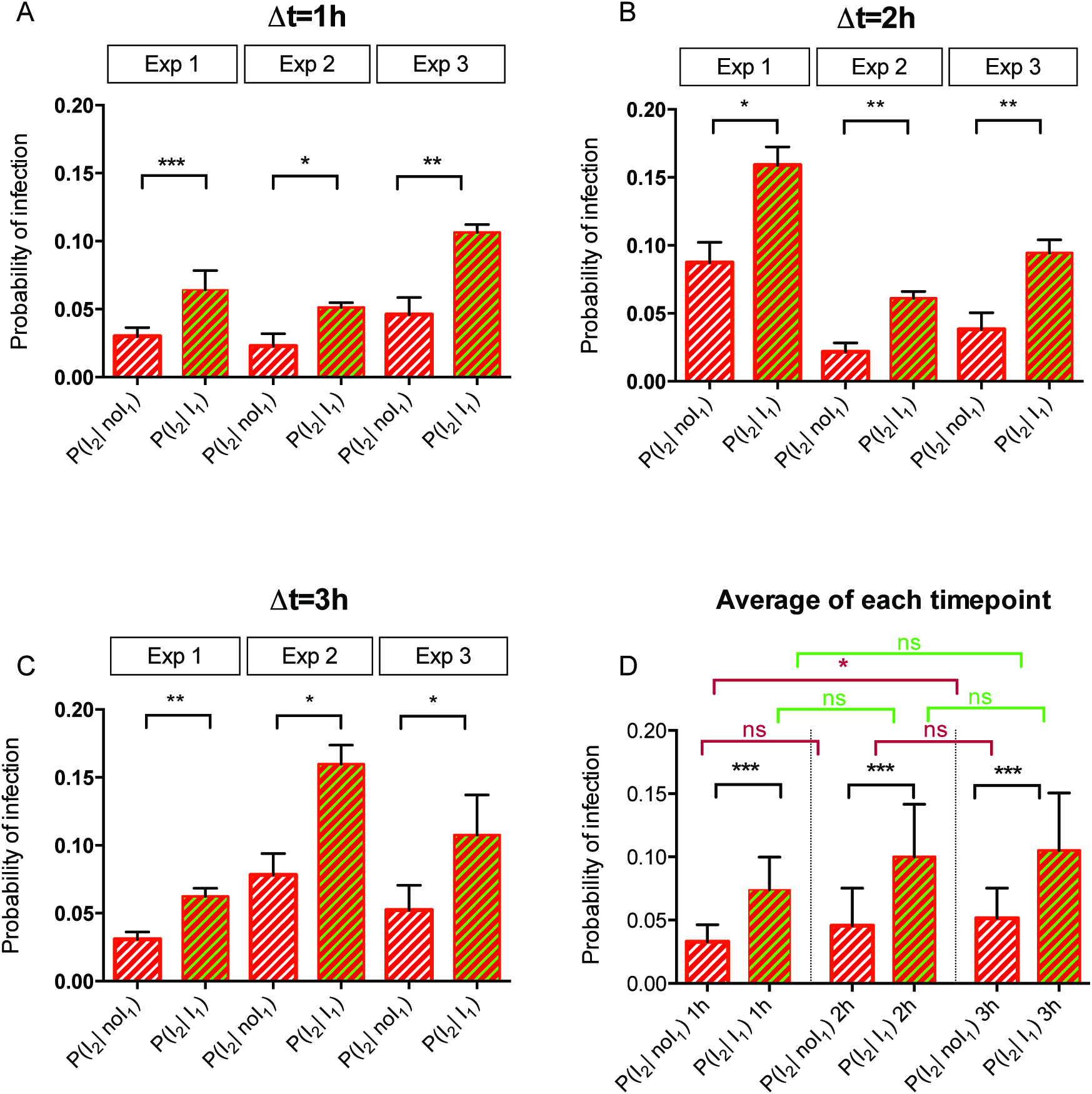
**A.B.C.** Detailed depiction of the conditional probability of infection for two different populations during sequential infection with a delay of 1 h (**A**), 2 h (**B**) and 3 h (**C**) for each independent experiment with 3 replicates per experiment. P-values were obtained after paired t-test. **D.** Representation of the results from **A**, **B** and **C** after averaging them for each delay. P-values were obtained after paired t-test. The P-values in black resulted from a t-test comparing P(I_2_ | I_1_) and P(I_2_| noI_1_). The P-values in red resulted from a t-test comparing P(I_2_| noI_1_) for 1 h versus 2 h and 2 h versus 3 h. The P-values in green resulted from a t-test comparing P(I_2_| I_1_) for 1 h versus 2 h and 2 h versus 3 h.

**Fig. S3.**
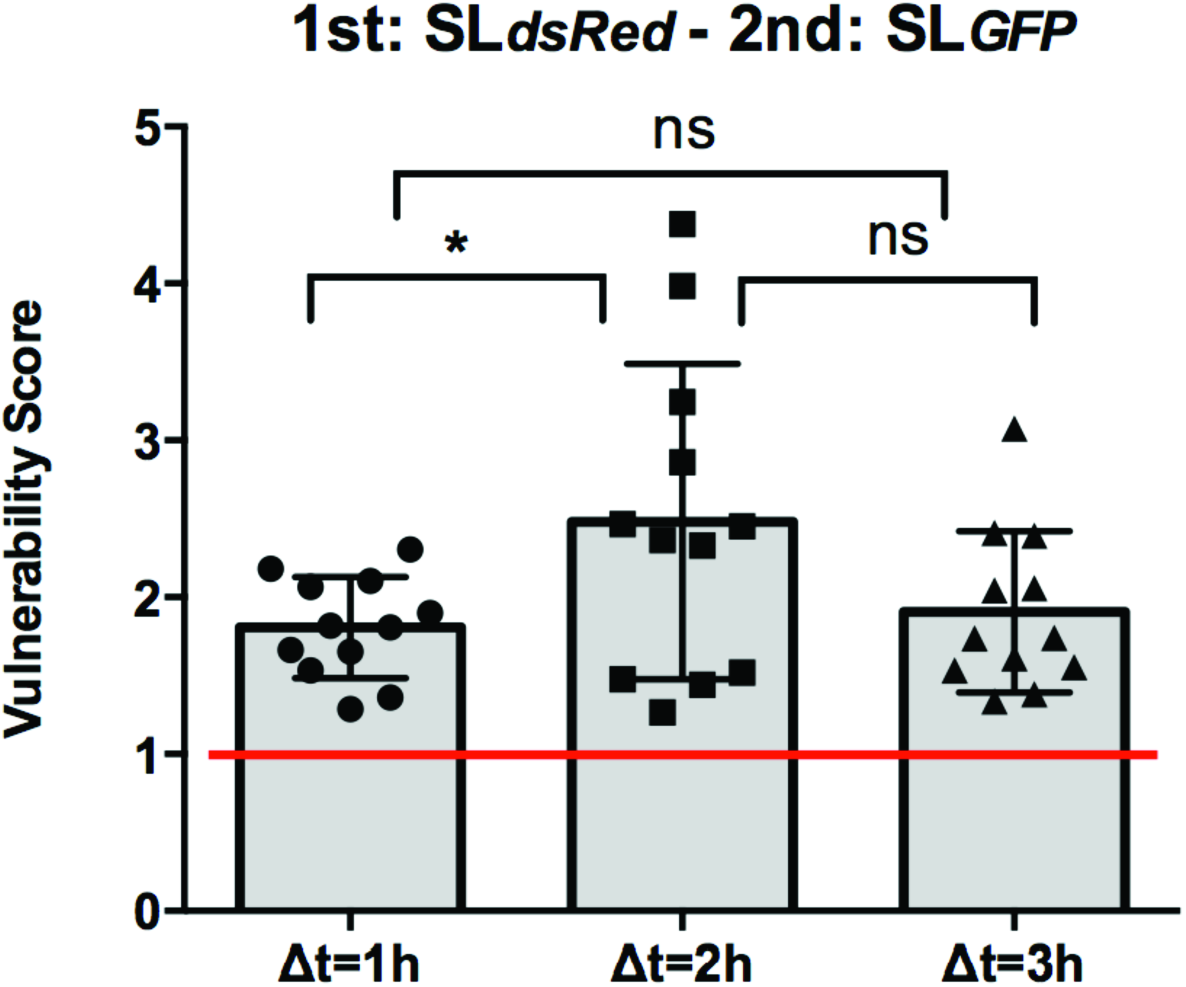
Vulnerability scores for the inverted infections compared to Fig.2C (SL*_dsRed_* before SL*_GFP_*) with a delay of 1, 2 and 3 h between infections. The red line corresponds to P(I_2_ | I_1_)=P(I_2_ | noI_1_) = 1 indicating the independence of the infections I_2_ and I_1_. Values above the red line correspond to P(I_2_ | I_1_) >P(I_2_ | noI_1_) indicating a cooperation between infections. Values below the red line correspond to P(I_2_ | I_1_) <P(I_2_ | noI_1_) indicating a competition between infections. Results were obtained from 3 independent experiments per time-point, and P-values were obtained after unpaired t-test.

**Fig. S4.**
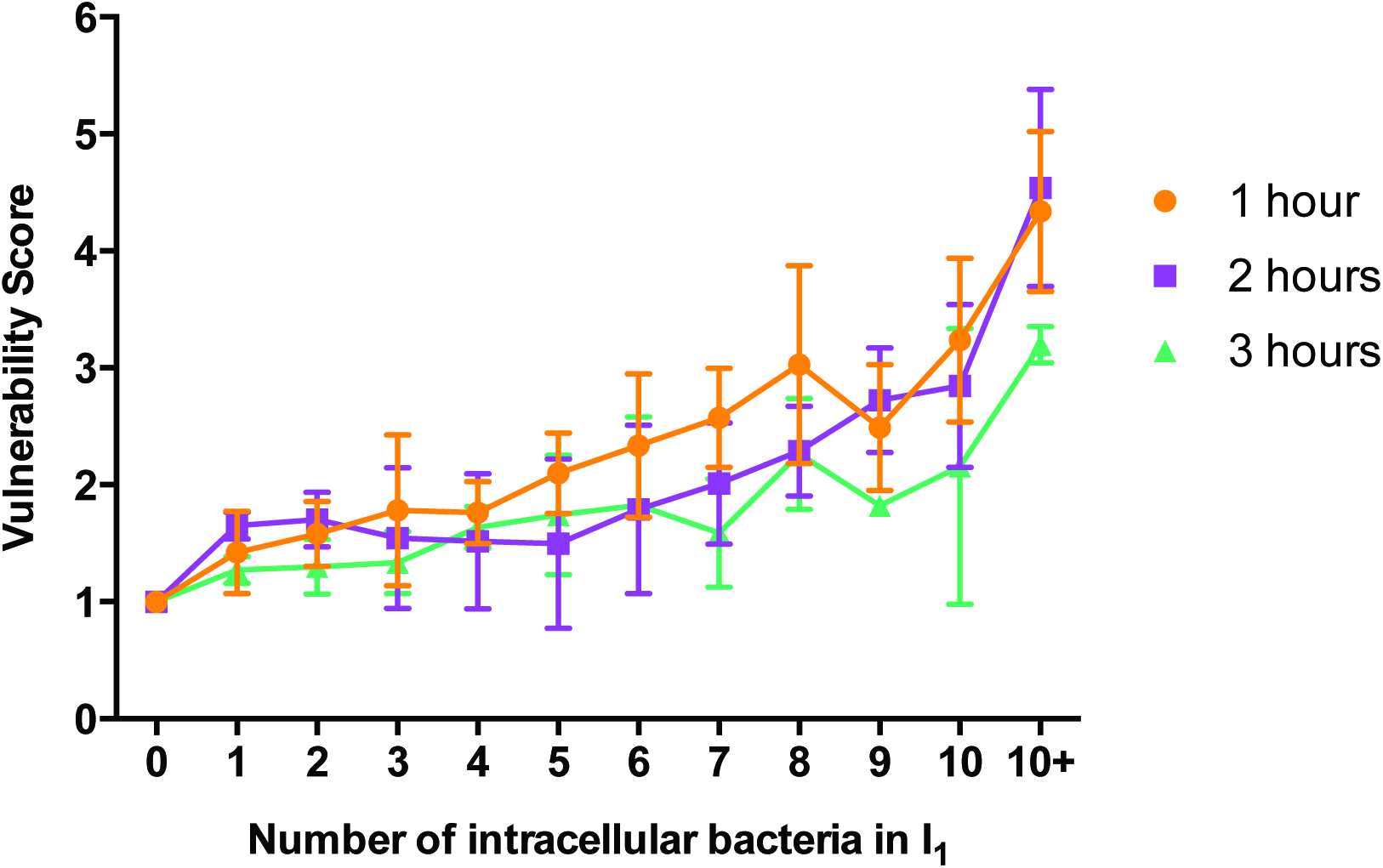
Vulnerability score as a function of the number of intracellular bacteria resulting from the 1^st^ infection with a delay of 1, 2 and 3 h between the infections. Results were obtained from 3 independent experiments per time-point.

**Fig. S5.**
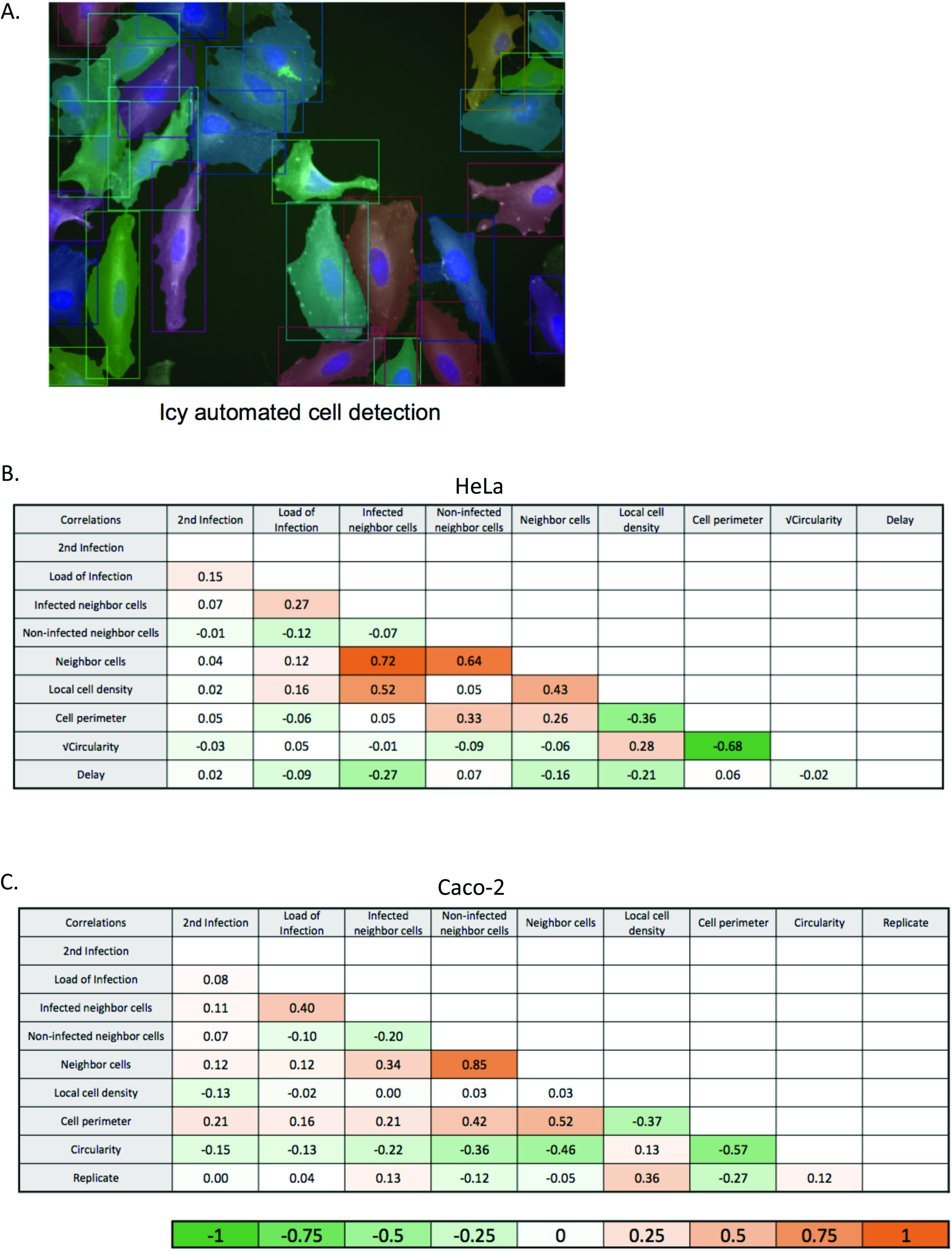
Cell parameter correlations. A. Illustration of Icy cell segmentation using *Active Contours* (*see Materials and Methods for plugins detail*). **B-C.** Table of the correlations between the different cell parameters for HeLa (**B**) and Caco-2 (**C**) cells.

**Fig. S6.**
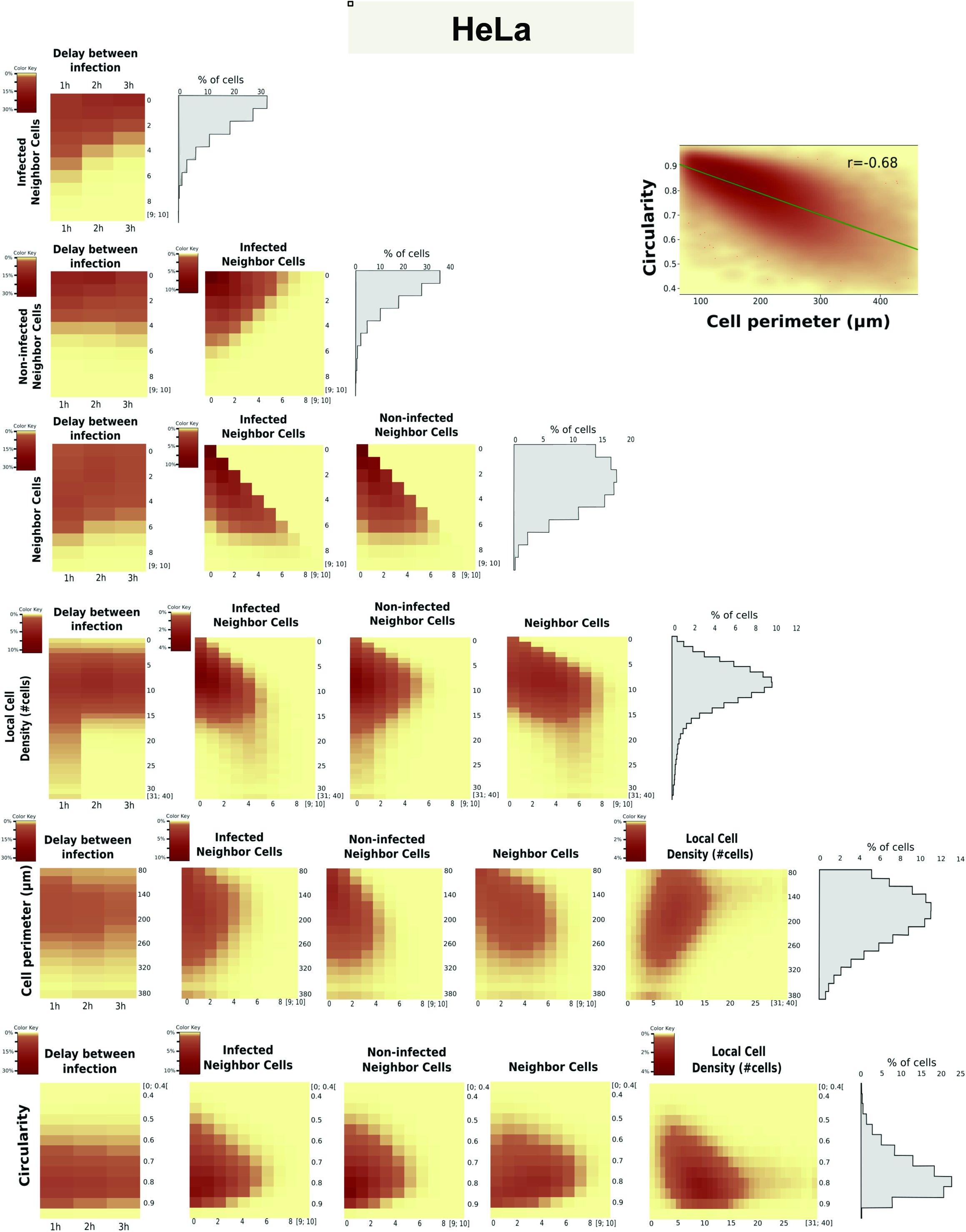
Scatter plot and heat-map of the different cell parameters studied in Hela cells model allowing to evaluate the relation between these parameters. Grey histograms represent the distribution of the vertical axe parameter in the entire cell population.

**Fig. S7.**
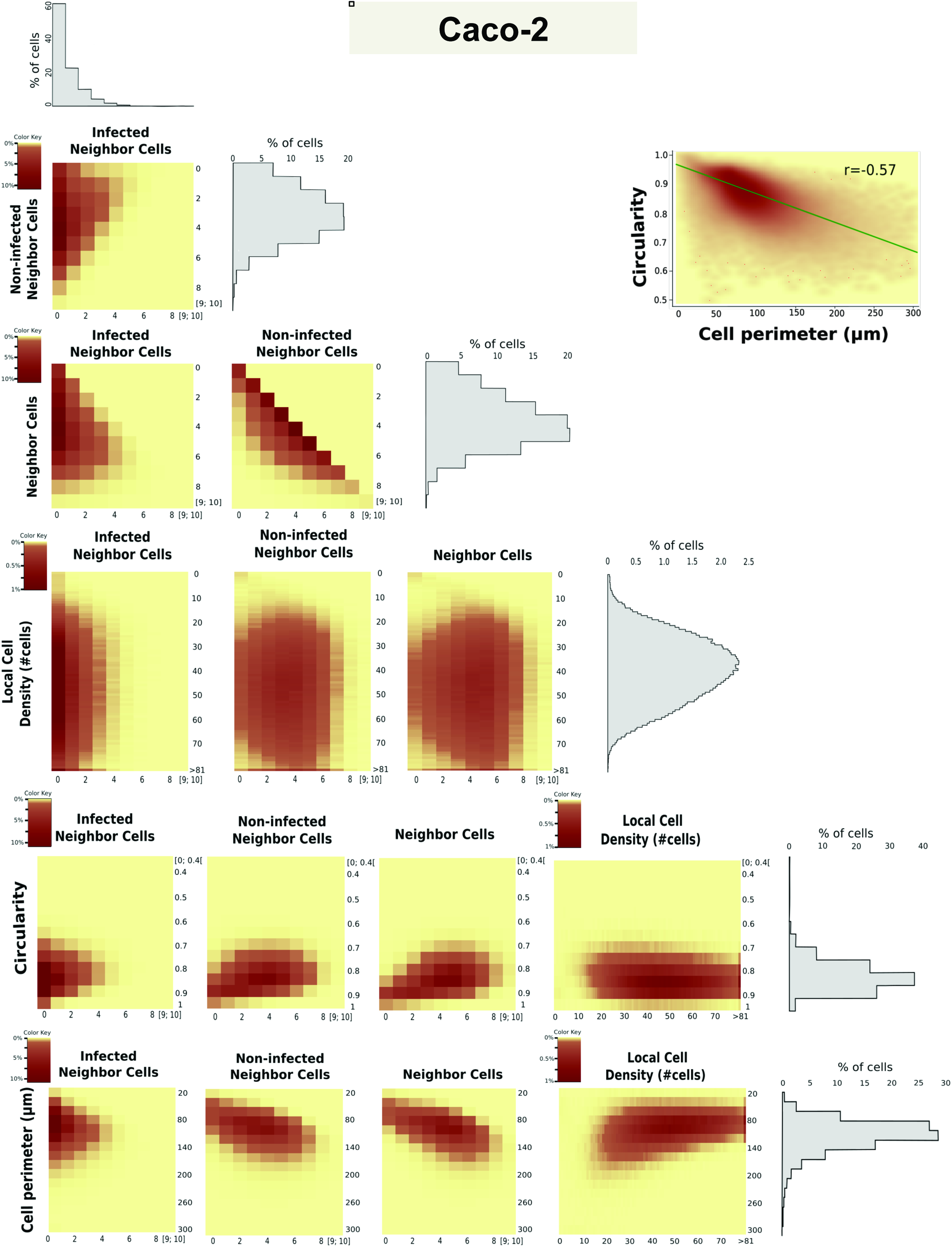
Scatter plot and heat-map of the different cell parameters studied in Caco-2 cells model allowing to evaluate the relation between these parameters. Grey histograms represent the distribution of the parameter in the entire cell population.

**Fig. S8.**
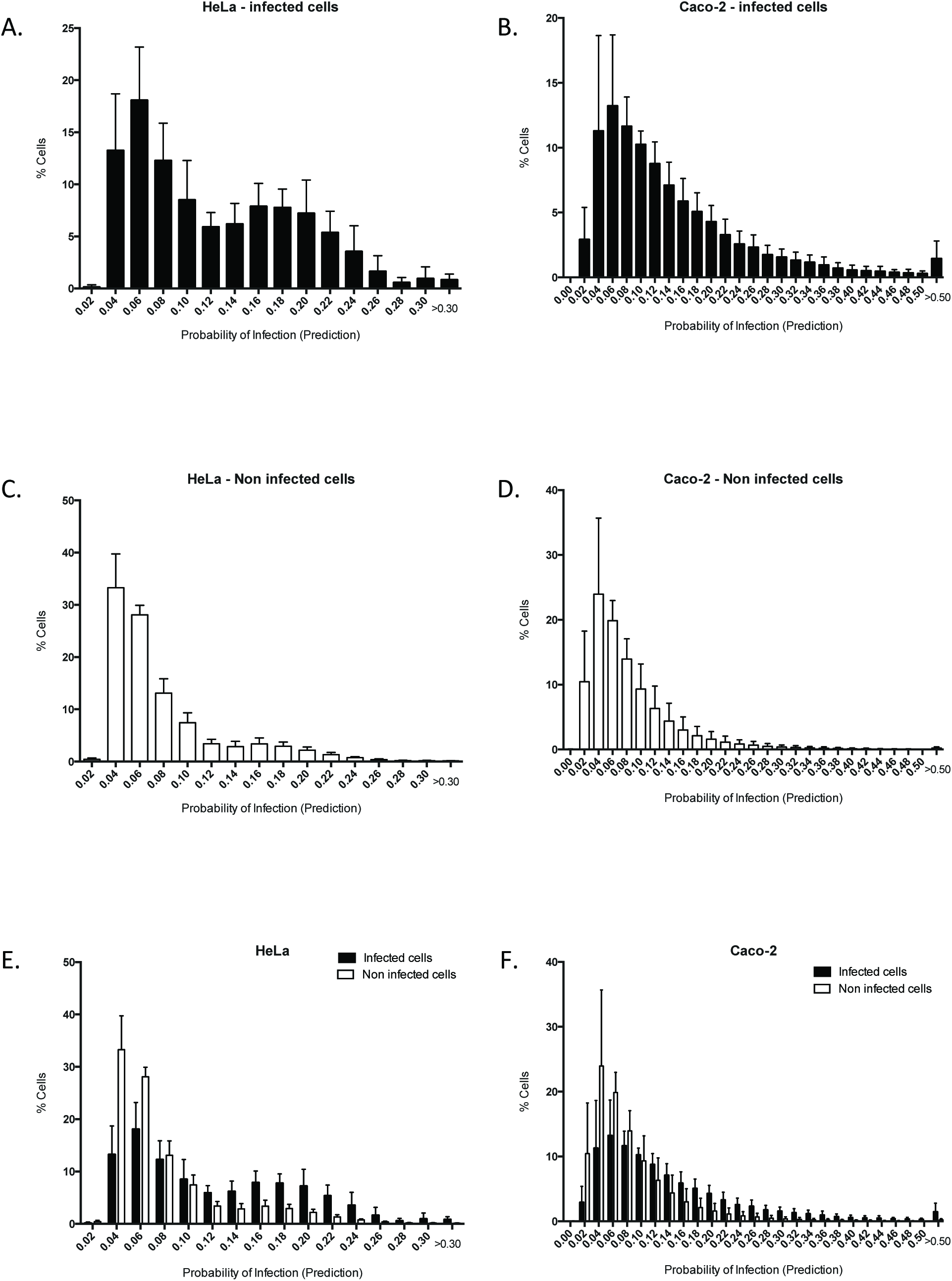
Distribution of the predicted probability of infection at single-cell level for infected and non infected cells. **A-C-E.** HeLa cells. **B-D-F.** Caco-2 cells.

**Fig. S9.**
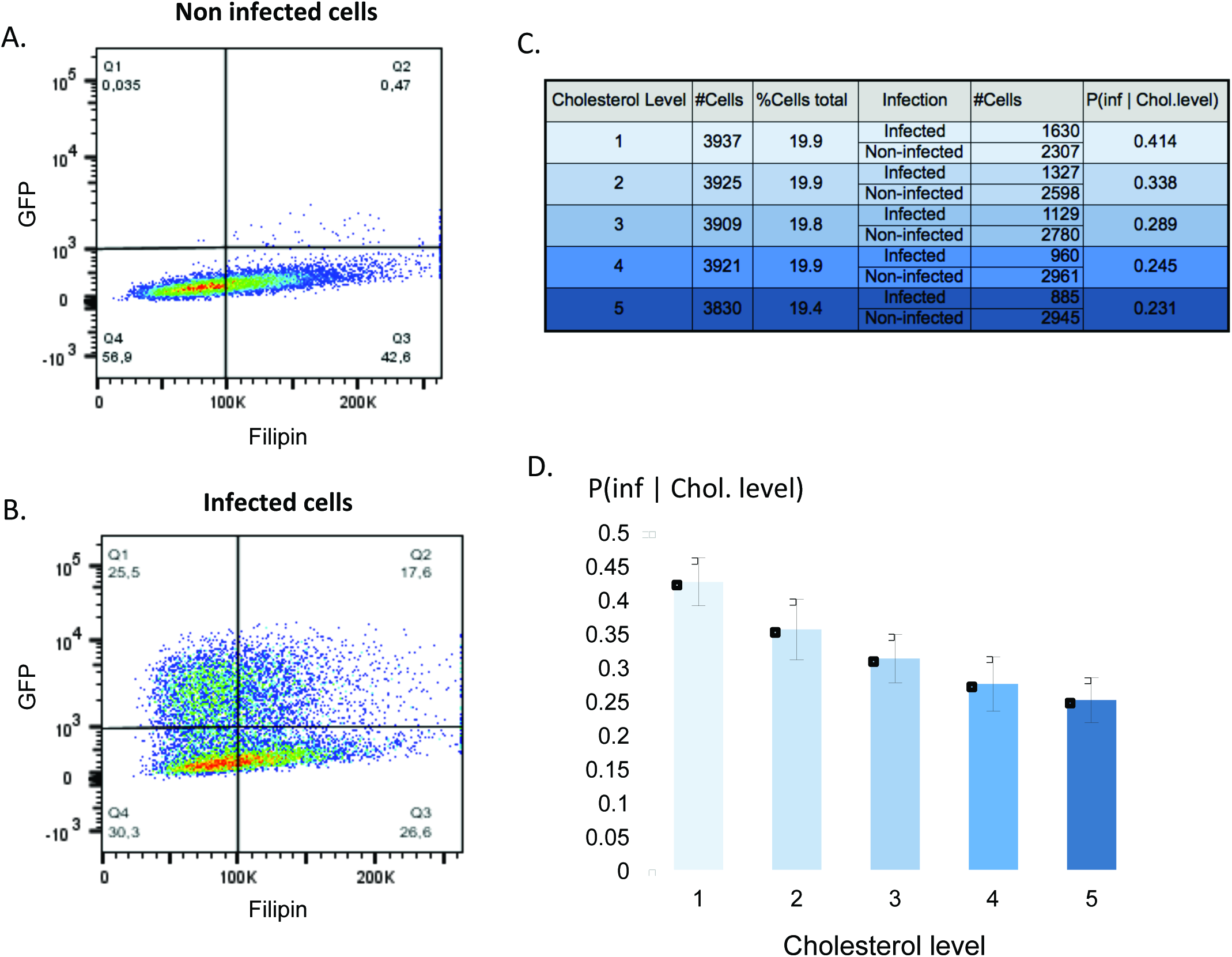
Illustration of FACS acquisition and data processing. **A-B.** Heat map of Filipin fluorescence as measure of the host cholesterol level (horizontal axe) and GFP fluorescence representing Salmonella-GFP infection (vertical axe). The dials [Q1,Q2] and [Q3,Q4] correspond to the non-infected (**A**) and the infected cells respectively (**B**). **C.** Raw data obtained after binning of the total cell population in 5 categories of cholesterol level (from the lowest to the highest) containing approximately the same number of cell (column 1 to 4). Conditional probability of infection for each category of cholesterol level based on the raw data (5^th^ column). **D.** Representation of the conditional probability of infection for each category of cholesterol level.

## REFERENCE

1. Carpenter, A. E. et al. CellProfiler: image analysis software for identifying and quantifying cell phenotypes. Genome Biol. 7, R100 (2006).

2. Carter, P. B. & Collins, F. M. The route of enteric infection in normal mice. J. Exp. Med. 139, 1189–203 (1974).

3. Criddle, A., Thornburg, T., Kochetkova, I., DePartee, M. & Taylor, M. P. gD-Independent Superinfection Exclusion of Alphaherpesviruses. J. Virol. 90, 4049–4058 (2016).

4. Cumby, N., Davidson, A. R. & Maxwell, K. L. The moron comes of age. Bacteriophage 2, 225–228 (2012).

5. Davey, M. E. & O'toole, G. A. Microbial biofilms: from ecology to molecular genetics. Microbiol. Mol. Biol. Rev. 64, 847–67 (2000).

6. de Chaumont, F. et al. Icy: an open bioimage informatics platform for extended reproducible research. Nat. Methods 9, 690–696 (2012).

7. Doceul, V., Hollinshead, M., van der Linden, L. & Smith, G. L. Repulsion of superinfecting virions: a mechanism for rapid virus spread. Science 327, 873–876 (2010).

8. Ehsani, S. et al. Hierarchies of host factor dynamics at the entry site of Shigella flexneri during host cell invasion. Infect. Immun. 80, 2548–2557 (2012).

9. Frechin, M. et al. Cell-intrinsic adaptation of lipid composition to local crowding drives social behaviour. Nature 523, 88–91 (2015).

10. Garner, M. J., Hayward, R. D. & Koronakis, V. The Salmonella pathogenicity island 1 secretion system directs cellular cholesterol redistribution during mammalian cell entry and intracellular trafficking. Cell. Microbiol. 4, 153–165 (2002).

11. Gilk, S. D. et al. Bacterial Colonization of Host Cells in the Absence of Cholesterol. PLoS Pathog. 9, (2013).

12. Haraga, A., Ohlson, M. B. & Miller, S. I. Salmonellae interplay with host cells. Nat. Rev. Microbiol. 6, 53–66 (2008).

13. Hayward, R. D. et al. Cholesterol binding by the bacterial type III translocon is essential for virulence effector delivery into mammalian cells. Mol. Microbiol. 56, 590–603 (2005).

14. Huang, I. C. et al. Influenza A virus neuraminidase limits viral superinfection. J Virol 82, 4834–4843 (2008).

15. Jones, B. D., Paterson, H. F., Hall, a & Falkow, S. Salmonella typhimurium induces membrane ruffling by a growth factor-receptor-independent mechanism. Proc. Natl. Acad. Sci. U. S. A. 90, 10390–10394 (1993).

16. Jorgensen, I. et al. The chlamydia protease CPAF regulates host and bacterial proteins to maintain pathogen vacuole integrity and promote virulence. Cell Host Microbe 10, 21–32 (2011).

17. Kamentsky, L. et al. Improved structure, function and compatibility for cellprofiler: Modular high-throughput image analysis software. Bioinformatics 27, 1179–1180 (2011).

18. Karpf, a R., Lenches, E., Strauss, E. G., Strauss, J. H. & Brown, D. T. Superinfection exclusion of alphaviruses in three mosquito cell lines persistently infected with Sindbis virus. J. Virol. 71, 7119–7123 (1997).

19. Kasper, C. A. et al. Cell-cell propagation of NF-??B transcription factor and MAP kinase activation amplifies innate immunity against bacterial infection. Immunity 33, 804–816 (2010).

20. Knodler, L. A. Salmonella enterica: Living a double life in epithelial cells. Curr. Opin. Microbiol. 23, 23–31 (2015).

21. Laliberte, J. P. & Moss, B. A novel mode of poxvirus superinfection exclusion that prevents fusion of the lipid bilayers of viral and cellular membranes. J. Virol. 88, 9751–68 (2014).

22. LaRock, D. L., Chaudhary, A. & Miller, S. I. Salmonellae interactions with host processes. Nat. Rev. Microbiol. 13, 191–205 (2015).

23. Lelouard, H. et al. Pathogenic Bacteria and Dead Cells Are Internalized by a Unique Subset of Peyer's Patch Dendritic Cells That Express Lysozyme. Gastroenterology 138, 173–184.e3 (2010).

24. Liberali, P., Snijder, B. & Pelkmans, L. Single-cell and multivariate approaches in genetic perturbation screens. Nat. Rev. Genet. 16, 18–32 (2014).

25. Lorkowski, M., Felipe-López, A., Danzer, C. A., Hansmeier, N. & Hensel, M. Salmonella enterica invasion of polarized epithelial cells is a highly cooperative effort. Infect. Immun. 82, 2657–2667 (2014).

26. Majowicz, S. E. et al. The Global Burden of Nontyphoidal Salmonella Gastroenteritis. Clin. Infect. Dis. 50, 882–889 (2010).

27. McQuate, S. E. et al. Long-term live-cell imaging reveals new roles for *Salmonella* effector proteins SseG and SteA. Cell. Microbiol. (2016). doi:10.1111/cmi.12641

28. Misselwitz, B. et al. Near surface swimming of salmonella Typhimurium explains target-site selection and cooperative invasion. PLoS Pathog. 8, 9 (2012).

29. Misselwitz, B. et al. Salmonella enterica serovar typhimurium binds to hela cells via fim-mediated reversible adhesion and irreversible type three secretion system 1-mediated docking. Infect. Immun. 79, 330–341 (2011).

30. Pace, J., Hayman, M. J. & Galán, J. E. Signal transduction and invasion of epithelial cells by S. typhimurium. Cell 72, 505–514 (1993).

31. Pier, G. B. et al. Salmonella typhi uses CFTR to enter intestinal epithelial cells. Nature 393, 79–82 (1998).

32. Santos, A. J. M., Meinecke, M., Fessler, M. B., Holden, D. W. & Boucrot, E. Preferential invasion of mitotic cells by Salmonella reveals that cell surface cholesterol is maximal during metaphase. J. Cell Sci. 126, 2990–2996 (2013).

33. Santos, J. C. & Enninga, J. At the crossroads: Communication of bacteria-containing vacuoles with host organelles. Cell. Microbiol. 18, 330–339 (2016).

34. Schaller, T. et al. Analysis of hepatitis C virus superinfection exclusion by using novel fluorochrome gene-tagged viral genomes. J. Virol. 81, 4591–603 (2007).

35. Snijder, B. et al. Population context determines cell-to-cell variability in endocytosis and virus infection. Nature 461, 520–523 (2009).

36. Stecher, B. et al. Flagella and Chemotaxis Are Required for Efficient Induction of Salmonella enterica Serovar Typhimurium Colitis in Streptomycin-Pretreated Mice Flagella and Chemotaxis Are Required for Efficient Induction of Salmonella enterica Serovar Typhimurium Colitis. Infect Immun 72, 4138–4150 (2004).

37. Vonaesch, P. et al. Quantitative insights into actin rearrangements and bacterial target site selection from SalmonellaTyphimurium infection of micropatterned cells. Cell. Microbiol. 15, 1851–1865 (2013).

38. Watson, K. G. & Holden, D. W. Dynamics of growth and dissemination of Salmonella in vivo. Cell. Microbiol. 12, 1389–1397 (2010).

39. Zou, G. et al. Exclusion of West Nile virus superinfection through RNA replication. J. Virol. 83, 11765–11776 (2009).

